# Dichotomous regulation of lysosomes by MYC and TFEB controls hematopoietic stem cell fate

**DOI:** 10.1101/2021.02.24.432720

**Authors:** Laura García-Prat, Kerstin B. Kaufmann, Florin Schneiter, Veronique Voisin, Alex Murison, Jocelyn Chen, Michelle Chan-Seng-Yue, Olga I. Gan, Jessica L. McLeod, Sabrina A. Smith, Michelle C. Shoong, Darrien Paris, Kristele Pan, Andy G.X. Zeng, Gabriela Krivdova, Kinam Gupta, Shin-Ichiro Takayanagi, Elvin Wagenblast, Weijia Wang, Mathieu Lupien, Timm Schroeder, Stephanie Z. Xie, John E. Dick

## Abstract

It is critical to understand how quiescent long-term hematopoietic stem cells (LT-HSC) sense demand from daily and stress-mediated cues and transition into bioenergetically active progeny to differentiate and meet these cellular needs. Here, we show that lysosomes, which are sophisticated nutrient sensing and signaling centers, are dichotomously regulated by the Transcription Factor EB (TFEB) and MYC to balance catabolic and anabolic processes required for activating LT-HSC and guiding their lineage fate. TFEB-mediated induction of the endolysosomal pathway causes membrane receptor degradation, limiting LT-HSC metabolic and mitogenic activation, which promotes quiescence, self-renewal and governs erythroid-myeloid commitment. By contrast, MYC engages biosynthetic processes while repressing lysosomal catabolism to drive LT-HSC activation. Collectively, our study identifies lysosomes as a central regulatory hub for proper and coordinated stem cell fate determination.

## Introduction

Human long-term hematopoietic stem cells (LT-HSC), at the apex of the hematopoietic hierarchy, must meet enormous daily demand (∼10^11^ cells daily) while also sustaining life-long maintenance of the stem cell pool. This hierarchical organization is widely thought to protect LT-HSC from exhaustion by their maintenance in a quiescent and undifferentiated state, activating only in response to microenvironment signals to generate highly proliferative but more short-lived populations including short-term HSC (ST-HSC) and committed progenitors(*1*). Upon cues to exit this dormant state, HSC must respond and adapt their metabolism and nutrient uptake to meet increased bioenergetic demands for cell growth and differentiation(*2*). Simultaneously, the events underlying cellular and metabolic activation must also be suppressed within a subset of LT-HSC to enable re-entry to quiescence and ultimately maintaining the LT-HSC pool through self-renewal(*2, 3*). However, the demand-adapted regulatory circuits of these early steps of hematopoiesis are largely unknown. Sensing signals or nutrient uptake depends on proteins that are embedded within the plasma membrane. These proteins internalize through endocytosis and can be degraded in the lysosomes or rerouted back to the cell surface and reused(*4*). Lysosomes are also terminal catabolic stations for autophagy where they clear cytoplasmic components, a process essential for preserving adult stem cell function(*5–9*). Recent work shows that lysosomes are not merely degradation stations, but also serve as major signaling centers for molecular complex assembly including mTORC1, AMPK, GSK3 and the inflammasome(*10*). These signaling complexes sense, integrate and facilitate cross-talk between diverse signals, and ultimately enable responses including autophagy, cell growth, membrane repair and microbe clearance. Although these distinct lysosomal activities in the stem cell context are largely unknown, lysosomes in mouse HSC were recently reported to be asymmetrically inherited, predicting future daughter cell fates(*11*), while pharmacological inhibition of the lysosomal v-ATPase positively impacted mouse HSC engraftment(*12*). Thus, we hypothesize that lysosomes coordinate the cell cycle and metabolic machinery of LT-HSC through their ability to sense and respond to diverse signaling cues to adapt their fate and lineage choices.

The MiT/TFE family of transcription factors (TFs) control lysosomes, with most studies focusing on TFEB as a master regulator of lysosomes. TFEB can sense and respond to stress signals and metabolic cues, including nutrient starvation or mitochondrial damage, by transcriptional activation of endocytosis, autophagy and lysosomal biogenesis genes(*13*). TFEB and MYC appear to compete for binding to the same chromatin regions in HeLa cells due to a high degree of binding sequence homology(*14*), but the biological relevance of this potential competition in a stem cell context is not understood. MYC regulates many aspects of metabolism, and plays a role in murine HSC by balancing self-renewal and differentiation(*15, 16*). However, the role of MYC in human HSC is unknown, and no previous studies of the role of TFEB in either human or mouse HSC function have been undertaken. Here, we uncovered a MYC-TFEB-mediated dichotomous regulation of lysosomal activity that is required to balance anabolic and catabolic processes that ultimately impact human LT-HSC fate determination.

## Results

### MYC drives LT-HSC activation by promoting anabolism and inhibiting lysosomal activity

During mitogenic activation from quiescence, LT-HSC enter a transitional state where transcriptional and metabolic changes occur to promote G0 exit with resultant entry to cell cycle. RNA-sequencing (RNA-seq) analysis of uncultured (quiescent, qLT-HSC) vs cultured (activated, aLT-HSC) LT-HSC sorted from human cord blood (hCB) showed that genes associated with biosynthetic processes, such as ribosome and mitochondria biogenesis, were the most enriched gene sets in aLT-HSC (**Fig. 1A;Fig. S1A**). This included *MYC* and known MYC targets that are involved in promoting cell cycle, protein synthesis and mitochondrial metabolism (**Fig. S1B;Fig. 1A,B;Table S1**). Indeed, concordant with mouse HSC(*17*), MYC protein abundance was barely detectable in qLT-HSC but greatly increased in aLT-HSC at 24 and 48h of *in vitro* culture, when cells are growing in size while still remaining undivided(*18*) (**Fig. 1C;Fig. S1C**). Conversely, autophagy-lysosomal pathway gene sets were among the most upregulated in qLT-HSC (**Fig. 1A;Fig. S1B;Table S1**). Interestingly, *TFEB* gene expression was enriched in qLT-HSC. Additionally TFEB, a protein typically only showing nuclear localization upon stress(*19*), was found to be nuclear in qLT-HSC (**Fig. 1B,C;Fig. S1D)**. By contrast, TFEB levels and nuclear localization were reduced in aLT-HSC (**Fig. 1C)**. We noted that other MiT/TFE family members were not expressed with the same trajectories across the human hematopoietic hierarchy(*20, 21*) (**Fig. S1E,F; Fig. S2A,B)**. Importantly, of the two MiT/TFE family members that bind to genetic elements of the CLEAR network (TFEB and TFE3)(*22*) and show nuclear localization, only TFEB changes to cytoplasmic localization upon activation (**Fig. S2C-E)**. MYC and TFEB expression anticorrelated at the single-cell level (**Fig. 1D**). LAMP1 staining revealed that aLT-HSC had expanded lysosomal mass compared to qLT-HSC (**Fig. 1C**). However, TFEB and genes involved in lysosome activity were found to be downregulated during LT-HSC activation suggesting this increased lysosomal content may be due to decreased lysosomal turnover (*23, 24*). We validated this idea by blocking lysosome acidification and degradation with short-term Bafilomycin (BAF) treatment and finding that qLT-HSC had higher accumulation of lysosomes compared to aLT-HSC (**Fig. 1E;Fig S3A)**. In agreement, aLT-HSC displayed a gradual decrease in lysosomal acidification indicating less functional lysosomes (**Fig S3B)**.

**Fig. 1.**
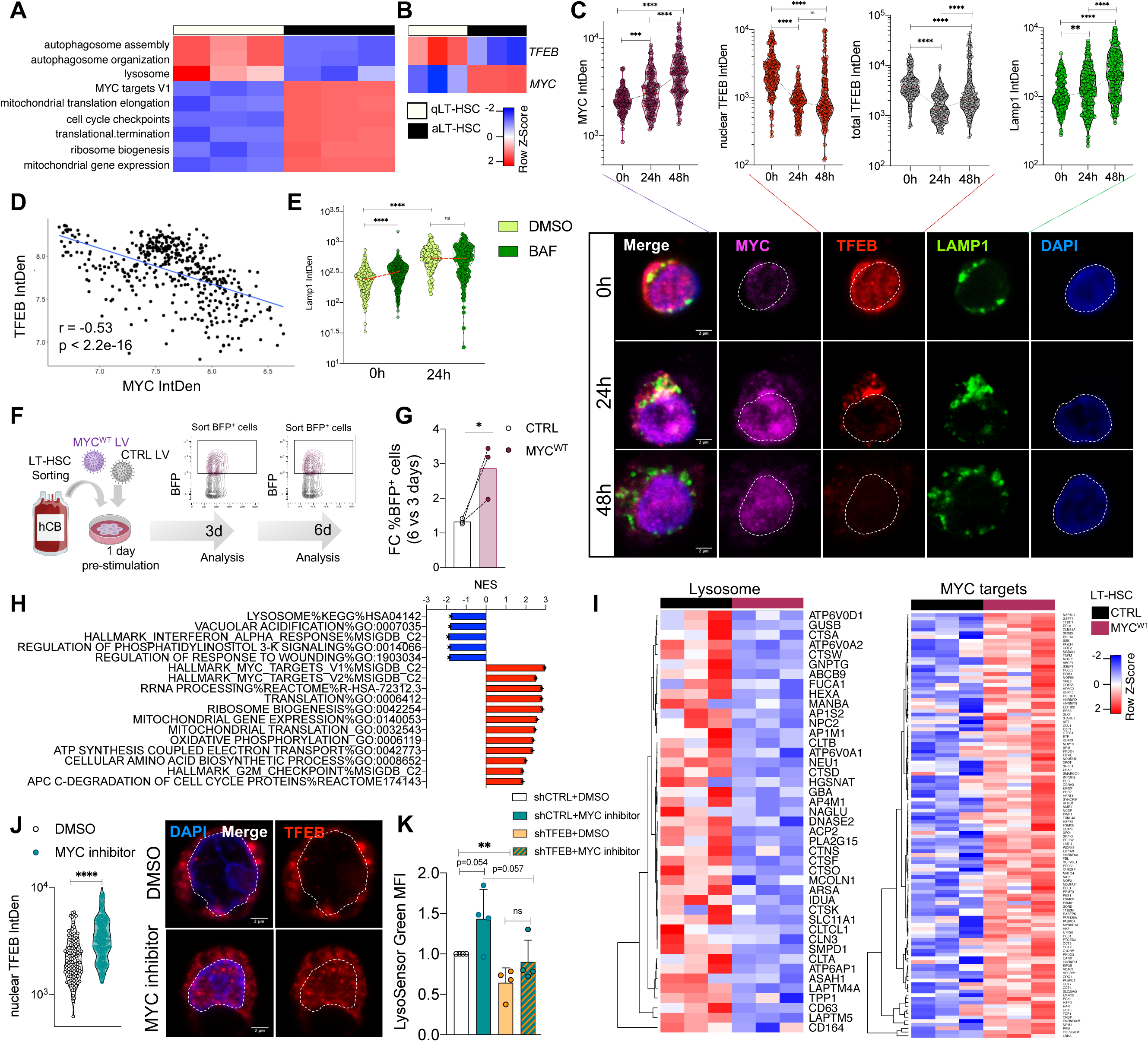
MYC drives LT-HSC activation by promoting anabolism and inhibiting lysosomal activity. **A.** RNA-seq analysis of qLT-HSC vs cultured LT-HSC for 4 days (d) (aLT-HSC). n=3CB. **B**. *MYC* and *TFEB* expression in LT-HSC from (A). **C**. Confocal analysis of TFEB, MYC and LAMP1 in qLT-HSC and cultured LT-HSC for 24h or 48h. IntDen (Integrated Density). Scale 2 µm. n=3CB, 469-867 cells/staining. Mann-Whitney test. **D**. Correlation of normalized IntDen values for MYC and TFEB from (C). **E**. Confocal analysis of LAMP1 in quiescent or 24h- cultured LT-HSC treated with DMSO or BAF for 3h. Mann-Whitney test. n=3CB, 594 cells. **F.** Scheme for LT-HSC transduced with lentivirus expressing BFP (blue fluorescent protein) and CTRL or MYC^WT^ genes. BFP^+^ cells were sorted at 3 or 6d for analysis. **G.** % BFP^+^ LT-HSC from (F) quantified by flow cytometry at 3 and 6d post-transduction. n=3CB. **H**. RNA-seq analysis of LT-HSC from (F) at 6d post-transduction. Normalized enrichment score (NES) of pathways differentially enriched in MYC^WT^ vs CTRL LT-HSC by GSEA analysis. FDRq- value<=0.05. n=3CB. **I**. Expression of indicated genes in LT-HSC from (F). **J.** Confocal analysis of TFEB in LT-HSC treated for 4d with DMSO or MYC inhibitor. n=3CB, 272 cells. Mann-Whitney test. **K.** LysoSensor mean fluorescence intensity (MFI) of LT-HSC transduced with lentivirus expressing mCherry and a shRNA against Renilla (shCTRL) or TFEB (shTFEB) treated with DMSO or MYC inhibitor from 1 to 5d post-transduction. n=4CB. *P<0.05, **P<0.01, ***P<0.001. Unpaired t-test unless otherwise indicated.

To directly assess MYC function in LT-HSC activation, we transduced LT-HSC with a MYC overexpressing lentivirus (LV) (MYC^WT^-OE) (**Fig. 1F**). MYC^WT^-OE LT-HSC doubled ∼3 fold more than control cells and at 6 days post-transduction they exhibited increased mitochondrial mass and expression of anabolic genes involved in ribosome biogenesis, protein translation, mitochondrial metabolism and MYC target genes (**Fig. 1G-I;Fig. S3C-F;Table S2**). Conversely, 4 days of MYC inhibition(*17*) restrained LT-HSC activation as demonstrated by reduced mitochondrial mass, ROS production and more G0-quiescent cells (**Fig. S3G-H**). MYC^WT^-OE downregulated lysosomal-related genes, consistent with our observation that lysosomal degradation is reduced in aLT-HSC (**Fig. 1H,I**). Importantly, MYC inhibition increased TFEB nuclear localization and lysosomal activity, which was blunted by *TFEB* knockdown (shTFEB) (**Fig. 1J,K**). Collectively, our data show that MYC is upregulated upon *in vitro* mitogenic stimulation causing repression of TFEB-associated lysosomal programs thereby driving LT-HSC activation and anabolism.

### TFEB-induced lysosomal activity inhibits MYC-driven anabolic processes and promotes LT-HSC quiescence

To gain insight into how TFEB regulates the transition from quiescence to activation, we performed transcriptomic analysis of control (CTRL) and TFEB-overexpressing (TFEB^WT^-OE) LT-HSC (**Fig. 2A;Fig. S4A**). As expected for a transcriptional activator, 67 out of 68 differentially expressed genes were upregulated in TFEB^WT^-OE compared to control LT-HSC (**Fig. S4B;Table S3**). Lysosomal-related gene sets were the most significantly enriched, including 14 different V-ATPase subunits that are required for lysosome and endosome acidification as well as cargo degradation (**Fig. 2B,C;Fig. S4C;Table S3**). Moreover, endosome transit to lysosomes or receptor endocytosis genes (such as transferrin or insulin receptor) were concurrently upregulated, indicating an overall induction of the endolysosomal pathway (**Fig. 2B-D)**. ATAC-Seq profiling uncovered 116 sites uniquely acquired in TFEB^WT^-OE vs 438 sites uniquely acquired in control LT-HSC (**Table S4**). TF recognition motif analysis identified specific E-boxes recognized by the MITF family of TFs as the most enriched binding motif in open chromatin regions gained upon TFEB^WT^-OE, while no significantly enriched motifs were found in control (**Fig. S4D,E**). Genes whose promoters fell within these gained elements showed enriched expression in TFEB^WT^-OE LT-HSC but depletion in shTFEB LT-HSC relative to controls providing strong evidence that the transcriptional changes observed are directly mediated by TFEB binding events (**Fig. S4F**). As expected, TFEB^WT^-OE increased LT-HSC lysosomal activity compared to control as demonstrated by the higher number, acidification and turnover of lysosomes as well as higher Cathepsin expression and activity (**Fig. 2E,F;Fig. S4G,H**). Notably, membrane protein levels of transferrin receptor 1 (TfR1), which is required for iron-bound transferrin uptake through clathrin-mediated endocytosis, were significantly reduced in TFEB^WT^-OE cells, despite only mild changes in levels of *TFRC* mRNA (**Fig. 2G;Fig. S4I)**. Confocal analysis confirmed the reduction of TfR1 membrane levels upon TFEB-OE and demonstrated the existence of TfR1^+^/LAMP1^+^ late endosomes/lysosomes (**Fig. 2F;Fig. S4J**). BAF treatment, previously shown to block TfR1 trafficking to late endosomes/lysosomes(*25*), or *LAMP1* knockdown (shLAMP1) resulted in increased TfR1 membrane levels, which could not be rescued by TFEB^WT^-OE (**Fig. 2H;Fig. S4K,L)**. Thus, we can conclude that TFEB induces clearance of TfR1 from the membrane of LT-HSC through endolysosomal degradation.

**Fig. 2.**
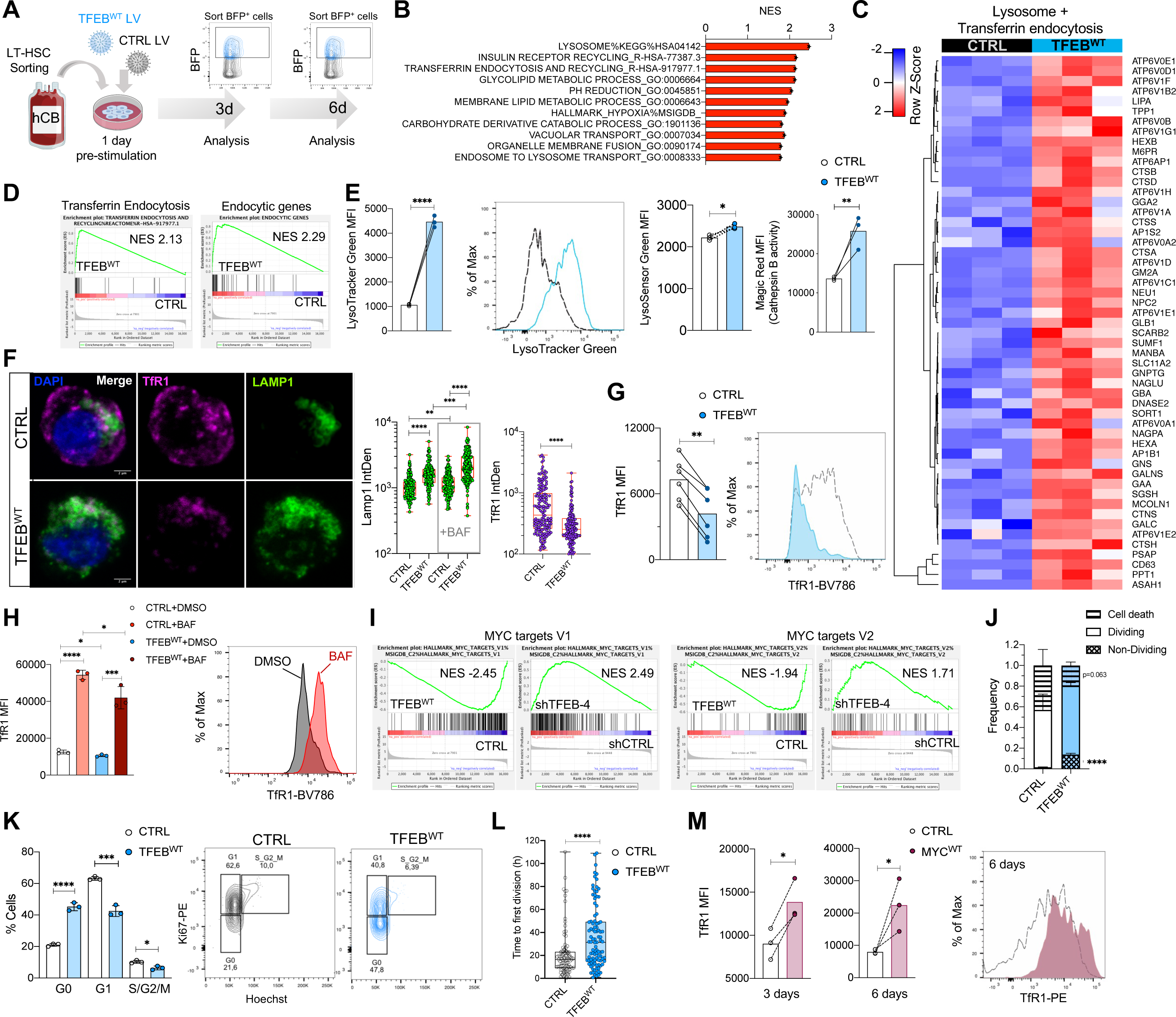
TFEB-induced endolysosomal degradation inhibits c-MYC-driven anabolic processes and promotes LT-HSC quiescence. **A.** Scheme for LT-HSC transduced with lentivirus expressing BFP and CTRL or TFEB^WT^ genes. BFP^+^ cells were sorted for analysis. **B.** RNA-seq analysis of LT-HSC from (A) at 3d post- transduction. NES value of pathways positively enriched in TFEB^WT^ vs CTRL LT-HSC by GSEA analysis. FDRq-value<=0.05. n=3CB. **C.** RNA-seq expression of lysosomal genes in LT- HSC from (A). **D.** GSEA plots of indicated gene sets in LT-HSC from (A). FDRq-value<=0.05. **E.** LT-HSC from (A) stained with LysoTracker, LysoSensor and Magic Red and analyzed by flow cytometry at 6d post-transduction. n=3CB. **F.** Confocal analysis of LAMP1 and TfR1 in LT-HSC from (A) treated for 3h with BAF or DMSO at 6d post-transduction. Scale 2 µm. n=3CB, 635 cells. Mann-Whitney test. **G.** TfR1 MFI analyzed by flow cytometry in LT-HSC from (A). Paired t-test. **H**. TfR1 MFI analyzed by flow cytometry in shCTRL and shTFEB LT- HSC treated with DMSO or BAF from 1 to 5d post-transduction. n=3CB. **I**. GSEA plots of indicated gene sets in LT-HSC from (A) and Fig. S5H. FDRq-value<=0.05. **J.** Distribution of cell fate by time-lapse imaging analysis of LT-HSC from (A), from 3 to 6d post-transduction. n=3CB. Mann-Whitney test. **K**. Cell cycle analysis of LT-HSC from (A) at 6d post- transduction. n=3CB. **L**. Time to first division analysis of LT-HSC from (A) from 3 to 6d post- transduction in hours (h). Mann-Whitney test. n=3CB. 192 cells. **M.** TfR1 MFI analyzed by flow cytometry in LT-HSC from Fig. 1F at 3 and 6d post-transduction. n=3CB. Paired t-test. *P<0.05, **P<0.01, ***P<0.001. Unpaired t-test unless otherwise indicated.

We hypothesized that TFEB-induced endolysosomal degradation of cell surface receptors such as TfR1 is required to maintain LT-HSC quiescence, and that inhibition of this pathway could lead to enhanced TfR1 membrane levels and LT-HSC activation. Whilst qLT-HSC were mostly negative for TfR1, increased TfR1 membrane expression correlated with *MYC* elevation and preceded upregulation of *TFRC* mRNA levels during LT-HSC activation (**Fig, S4M,N**). Notably, transcriptomic analysis showed a large and consistent downregulation of gene sets involved in cell cycle regulation, RNA processing/transport and ribosome biogenesis upon TFEB^WT^-OE (**Fig. 2I;Fig. S5A-C;Table S3**). This group of anabolic genes included *MYC* and MYC-target genes and its downregulation correlated with lower levels of mitochondria and ROS (**Fig. 2I;Fig. S5A-D)**, indicating that TFEB maintains LT-HSC in a low metabolic state during culture. TFEB^WT^-OE restricted LT-HSC division over 6 days of time-lapse imaging compared to controls and cells remained Ki67 and EdU negative in flow cytometry analyses (**Fig. 2J,K;Fig. S5E,F)**, indicating that TFEB^WT^-OE inhibits LT-HSC quiescence exit. Furthermore, single cell tracking of TFEB^WT^-OE LT-HSC showed delayed division kinetics among dividing cells relative to control, resulting in an overall reduced cellular expansion (**Fig. 2L;Fig. S5E;Movie S1)**. Of note, the induction of this quiescent/low metabolic state upon TFEB^WT^-OE also protected LT-HSC from cell death during *in vitro* culture (**Fig. 2J**).

In contrast to TFEB^WT^-OE, shTFEB resulted in decreased lysosomal and Cathepsin B activity and higher expression of anabolic pathways and the number of cycling cells (**Fig. 1K**;**Fig. 2I**;**Fig. S5B,C,G-I;Table S5**). Indeed, the transcriptomic programs upregulated by shTFEB and MYC^WT^-OE in LT-HSC were highly concordant (**Fig. S5C**). In agreement, genes upregulated by TFEB^WT^-OE and downregulated by MYC^WT^-OE were enriched in qLT-HSC and in single cell RNA-seq subsets of bone marrow hematopoietic stem and progenitor cells classified as more dormant (non-primed)(*7, 26–28*) (**Fig. S6A,B**). Moreover, LT-HSC exhibited increased cell surface levels of TfR1 without significant changes in *TFRC* mRNA levels upon MYC^WT^-OE, whereas MYC inhibition reduced surface expression of TfR1, consistent with the observed MYC-directed repression of the endolysosomal pathway (**Fig. 1I**;**Fig. 2M;Fig. S4I;Fig. S6C**). Importantly, *TFRC* knockdown (shTFRC) reduced Ki67^+^ LT-HSC during culture, indicating that TfR1 is instrumental for LT-HSC activation (**Fig. S6D,E**). Overall, our findings demonstrate that TFEB suppresses MYC activation and restrains a MYC-driven anabolic program while inducing endolysosomal degradation of membrane receptors such as TfR1, maintaining a quiescent, low-metabolic state in LT-HSC.

### Lysosomal activity governs LT-HSC lineage specification

As TfR1 is required for erythropoiesis(*29*), we next examined whether lysosomal activity differs among subpopulations of LT-HSC with distinct TfR1 membrane levels and whether this correlates with differing lineage commitment potential. LT-HSC, sorted on the basis of higher (Lyso^H^) lysosomal activity in culture were reminiscent of TFEB-OE LT-HSC as they displayed higher acidification and levels of Cathepsin B activity and lower TfR1 surface expression levels than LT-HSC with low activity (Lyso^L^) **(Fig. 2E-G;Fig. 3A;Fig. S6F-H)**. Colony-forming cell (CFC) assays showed cloning efficiency was significantly higher from Lyso^L^ than Lyso^H^ LT- HSC, predominantly due to increased BFU-E (burst-forming unit-erythroid), although total cellular output was unchanged **(Fig. 3B-D;Fig. S6I,J)**. The percentage and total number of cells expressing the erythroid marker GlyA and/or TfR1 were reduced in the progeny of Lyso^H^ compared to Lyso^L^ LT-HSC, while Lyso^H^ LT-HSC showed increased myeloid differentiation potential (CD33^+^/CD15^+^cells) **(Fig. 3C,D;Fig. S6J)**. Similarly, single-cell stromal (SCS) assays showed that myeloid and erythroid differentiation are associated with higher and lower lysosomal activity, respectively **Fig. S6K-N)**. Thus, the levels of lysosomal activity are inversely correlated with TfR1 surface expression levels and are tied to LT-HSC erythroid-myeloid lineage fate choices.

**Fig. 3.**
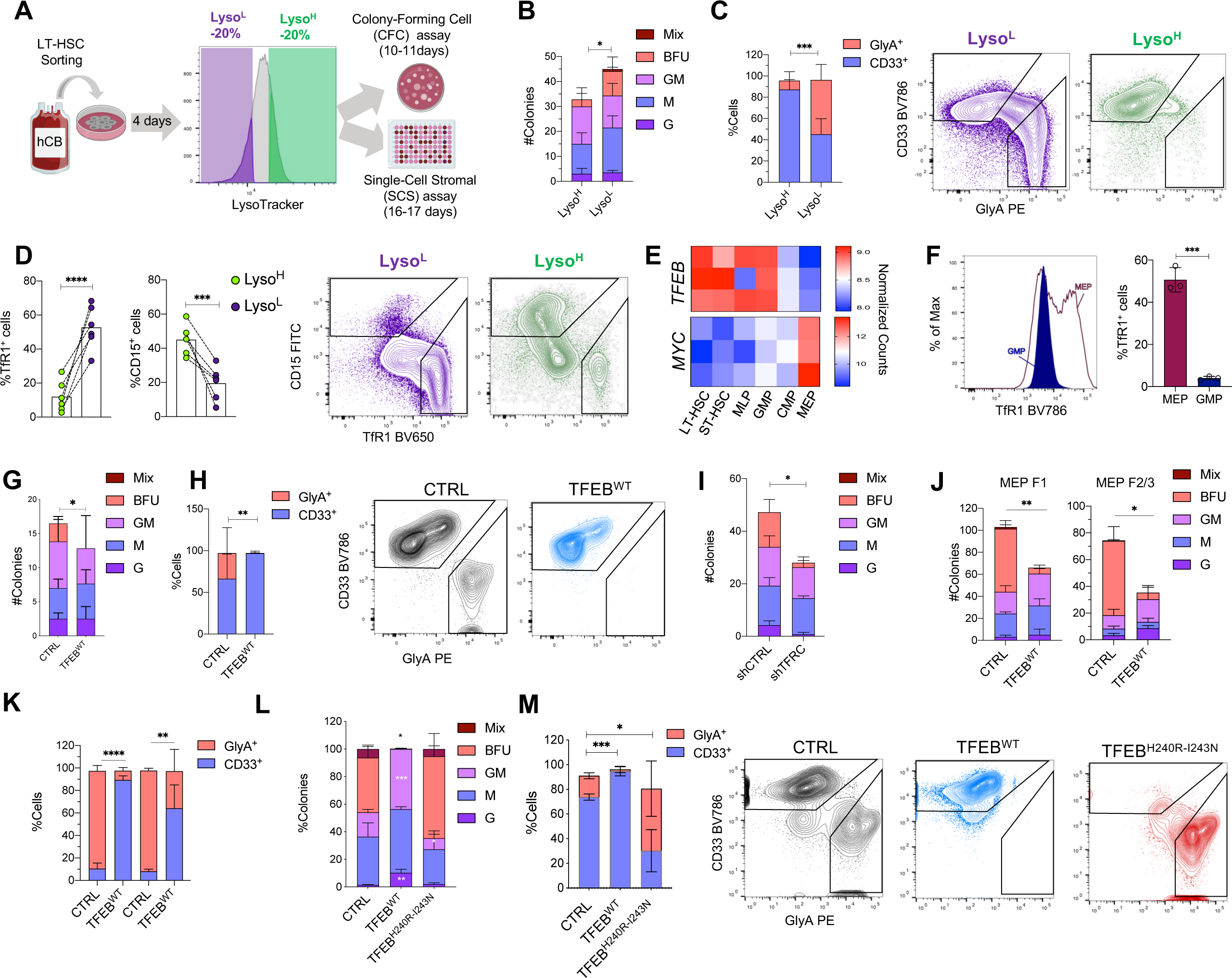
Lysosomal activity governs LT-HSC lineage specification. **A.** Experimental scheme: LT-HSC were cultured for 4d, stained with LysoTracker and sorted for the indicated differentiation assays. **B,C,D.** CFC colony distribution for LT-HSC from (A) scored under the microscope and GlyA+ and CD33+ lineage analysis by flow cytometry. n=3CB. **E.** RNA-seq normalized counts for *TFEB* and *MYC* in the indicated cell populations sorted as in Fig. S1A. n=3CB. **F.** TfR1 membrane expression analysis in MEP and GMP cells after 4d in culture. n=3CB. **G,H.** LT-HSC from Fig. S7C were plated for CFC assay as in (B,C), n=3CB. **I.** LT-HSC from Fig. S7R were plated for CFC assay as in (B,C), n=3CB. **J,K.** MEP F1 and MEPF2/3 cells from Fig. S7T were plated for CFC assay as in (B,C), n=3CB. **L,M.** CFC assay with CD34^+^CD38^-^ cells from Fig. S9B as in (B,C). n=4CB. *P<0.05, **P<0.01, ***P<0.001. Unpaired t-test unless otherwise indicated.

We asked whether similar regulation of TfR1-lysosomal degradation by TFEB and MYC might occur as part of the typical sequence of events that occurs during lineage commitment of LT-HSC. Interestingly, the gene expression patterns of *TFEB* and *MYC* in subpopulations downstream of LT-HSC were also mutually exclusive: megakaryocyte-erythrocyte progenitors (MEP) had low *TFEB* and high *MYC* expression, while granulocyte-monocyte progenitors (GMP) exhibited the inverse expression pattern **(Fig. 3E)**. TfR1 membrane levels were higher in MEP, correlating with *MYC* and anti-correlating with *TFEB* expression **(Fig. 3E,F).** Analysis of more mature populations revealed that *TFEB* was distinctly suppressed in erythroid progenitors **(Fig. S7A,B).**

To determine functionally whether the lysosomal program is instrumental in regulating erythroid/myeloid commitment of LT-HSC, we assessed LT-HSC differentiation potential after modulation of TFEB expression. TFEB^WT^-OE in LT-HSC caused a complete block of erythroid differentiation as demonstrated by an almost complete absence of BFU-E colonies and GlyA^+^ cells in both CFC and SCS assays; similar results occurred in ST-HSC upon TFEB^WT^-OE **(Fig. 3G,H;Fig. S7C-K)**. shTFEB or shLAMP1 in LT-HSC produced the opposite phenotype **(Fig. S7L-Q)**. Importantly, shTfR1 also reduced BFU colony formation similar to TFEB^WT^-OE in LT-HSC, demonstrating that TfR1 is important for early erythroid-commitment **(Fig. 3I;Fig. S7R,S)**. Remarkably, TFEB^WT^-OE in committed MEP subpopulations (F1, F2/F3)(*21*) was sufficient to suppress TfR1 membrane levels and abolish their erythroid potential while slightly increasing their myeloid output in both CFC and SCS assays (**Fig. S1A;Fig. 3J,K;Fig. S7T,V;Fig. S8A-I**). Transcriptomic analysis of TFEB^WT^-OE MEP F2/3 subsets confirmed that TFEB imposes a myeloid-associated gene program and inhibits the expression of key transcription factors essential for erythrocyte and megakaryocyte differentiation, such as GATA1 or RUNX1 (**Fig. S8J-N;Table S6**). Thus, TFEB expression is sufficient to alter erythroid-myeloid fate decisions even in a progenitor subpopulation committed to erythropoiesis and provides strong evidence that TFEB-mediated induction of the endolysosomal pathway controls early lineage determination in LT-HSC as well as downstream progenitors.

To determine whether TFEB may exert its functions through mechanisms other than transcriptional activation, we overexpressed either TFEB^WT^ or a TFEB variant unable to activate transcription due to disruption of its DNA binding domain (TFEB^H240R-I243N^) in hCB CD34^+^CD38^-^ cells (**Fig. S9A-D**). Flow cytometric analysis showed that overexpression of TFEB^WT^ but not TFEB^H240R-I243N^ increased autophagosome/lysosome formation and restrained expansion of CD34^+^CD38^-^ cells *in vitro* (**Fig. S9E,F**). In CFC assays, TFEB^WT^ strongly inhibited erythroid commitment and enhanced myeloid differentiation, whereas TFEB^H240R-I243N^ expression produced the opposite effect (**Fig. 3L,M**). These results indicate that TFEB exerts its functions primarily through transcriptional activation. We speculated that TFEB^H240R-I243N^ acts in a dominant negative manner, perhaps through formation of homodimers sequestering other TFEB molecules. These results independently support our findings with TFEB^WT^-OE and establish that TFEB-mediated induction of lysosomal activity is required for myelopoiesis but is incompatible with erythroid differentiation. Overall, our study has uncovered lysosomes as transcriptionally-controlled central signaling hubs that govern lineage determination.

### Enhanced TFEB-driven lysosomal activity distinguishes LT-HSC from ST-HSC

ST-HSC are the immediate downstream progeny of LT-HSC and share many transcriptional and functional properties. However, they differ in their quiescence exit kinetics and self-renewal properties(*18*). Since our data indicate that TFEB imposes a metabolically-low quiescent state in LT-HSC and restrains their expansion while inducing endolysosomal degradation of specific membrane receptors, we hypothesized that the TFEB-controlled lysosomal program could be differentially regulated in LT- vs ST-HSC. In relation to qST-HSC, qLT-HSC showed transcriptional enrichment of lysosomal pathway genes and contained variable but overall higher levels of lysosomes (*21, 30*) (**Fig. 4A,B;Fig. S9G,H;Table S7**). The observed enrichment of lysosomal genes was associated with higher levels of nuclear TFEB and lower levels of MYC in qLT- vs qST-HSC (**Fig. 4A**). In short term cultures, aLT-HSC also showed enrichment of *TFEB* and lysosomal genes with depletion of *MYC* and its target genes in comparison to aST-HSC (**Fig. 4C,D;Fig. S9I;Table S8**). Consistent with these transcriptomic results, aLT-HSC exhibited enhanced lysosomal activity as defined by Cathepsin B activity and reduced levels of mitochondria, ROS and TfR1 compared to aST-HSC (**Fig. 4E-H**). TFEB^WT^- OE in ST-HSC induced a transcriptional program similar to that of LT-HSC, with enrichment of endolysosomal genes and depletion of cell cycle and biosynthetic related genes (**Fig. 4I,J;Fig. S9J,K;Table S9**). Functionally, TFEB-OE in ST-HSC induced lysosomal activity with inhibition of quiescence exit and reduction in TfR1 membrane levels (**Fig. 4K-M;Fig. S9L**). MYC inhibition suppressed mitogenic and metabolic activation of ST-HSC in culture (**Fig. S9M**). Thus, aspects of a LT-HSC-specific program can be imposed on ST-HSC upon ectopic TFEB expression or MYC inhibition. Overall, these findings indicate that an enhanced TFEB-regulated lysosomal program distinguishes LT- from ST-HSC and actively restrains the anabolic processes of LT-HSC with resultant limitation in their cellular output upon mitogenic activation.

**Fig. 4.**
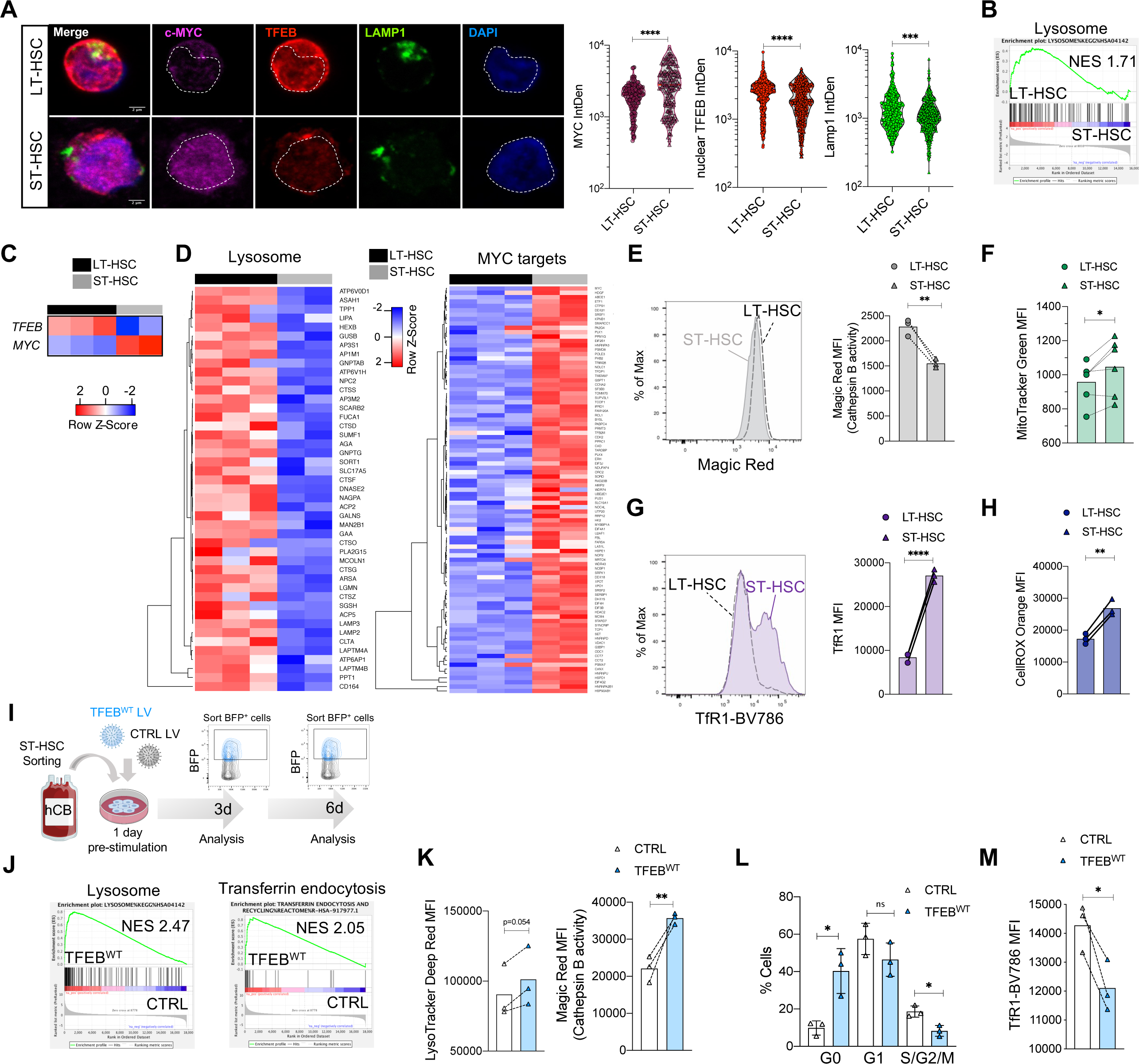
Enhanced TFEB-driven lysosomal activity differentiates LT from ST-HSC. **A,B.** Confocal analysis of LT-HSC and ST-HSC stained for TFEB, LAMP1, MYC and DAPI. Scale 2 µm. n=3CB, 239-951 individual cells/staining. Mann-Whitney test. **C,D.** RNA-seq expression of indicated genes in LT- and ST-HSC cultured for 4d. n=2-3CB. **E**,**F,G,H**. Magic Red, TfR1, MitoTracker and CellROX MFI in LT- and ST-HSC cultured for 4d and analyzed by flow cytometry. n=3-6CB. Paired t-test. **I**. Experimental scheme: ST-HSC transduced with lentiviral vectors expressing BFP and CTRL or TFEB^WT^ genes. BFP^+^ cells were sorted at 3 and 6d for the indicated analyses. **J.** Expression of indicated genes in ST-HSC from (I). n=2CB. **K**. LysoTracker and Magic Red analyzed by flow cytometry in ST-HSC from (I). n=3CB. **L.** Cell cycle analysis of ST-HSC from (I). n=3CB. **M.** TfR1 MFI analyzed by flow cytometry on ST-HSC from (I). n=3CB. *P<0.05, **P<0.01, ***P<0.001. Unpaired t-test unless otherwise indicated.

### TFEB-regulated lysosomal activity controls LT-HSC self-renewal

We next investigated whether enhanced lysosomal activity in LT-HSC impacted on self-renewal, the hallmark trait of stem cells. TFEB^WT^-OE in LT-HSC resulted in an initial decrease but subsequent increase in total colonies and cells in serial re-plating CFC assays (**Fig. S10A-D)**. In contrast, shTFEB blunted LT-HSC self-renewal ability in secondary CFC assays, prompting us to directly assess LT-HSC self-renewal capacity using gold standard xenograft assays (**Fig. S10E,F**). CD34^+^CD38^-^ hCB cells transduced with a TFEB^WT^-OE lentiviral vector generated significantly smaller grafts in both NSG and NSG-W41 mice at 4 and 17 weeks compared to controls, in keeping with the restraint in total cellular output observed *in vitro* (**Fig. 5A,B;Fig. S10G-K**). By contrast, TFEB^H240R-I243N^-OE where mice showed only modest reductions (**Fig. 5A,B;Fig. S10G-K**). Remarkably, despite their lower repopulation TFEB^WT^, but not TFEB^H240R- I243N^ cells, showed significant enrichment of self-renewing LT-HSC as measured in serial xenotransplantation limiting dilution assays (LDA), with a 6.6 fold increase in stem cell frequency (**Fig. 5C;Fig. S10L,M**). These results suggest that TFEB governs LT-HSC self-renewal and exerts its effects primarily through DNA binding.

**Fig. 5.**
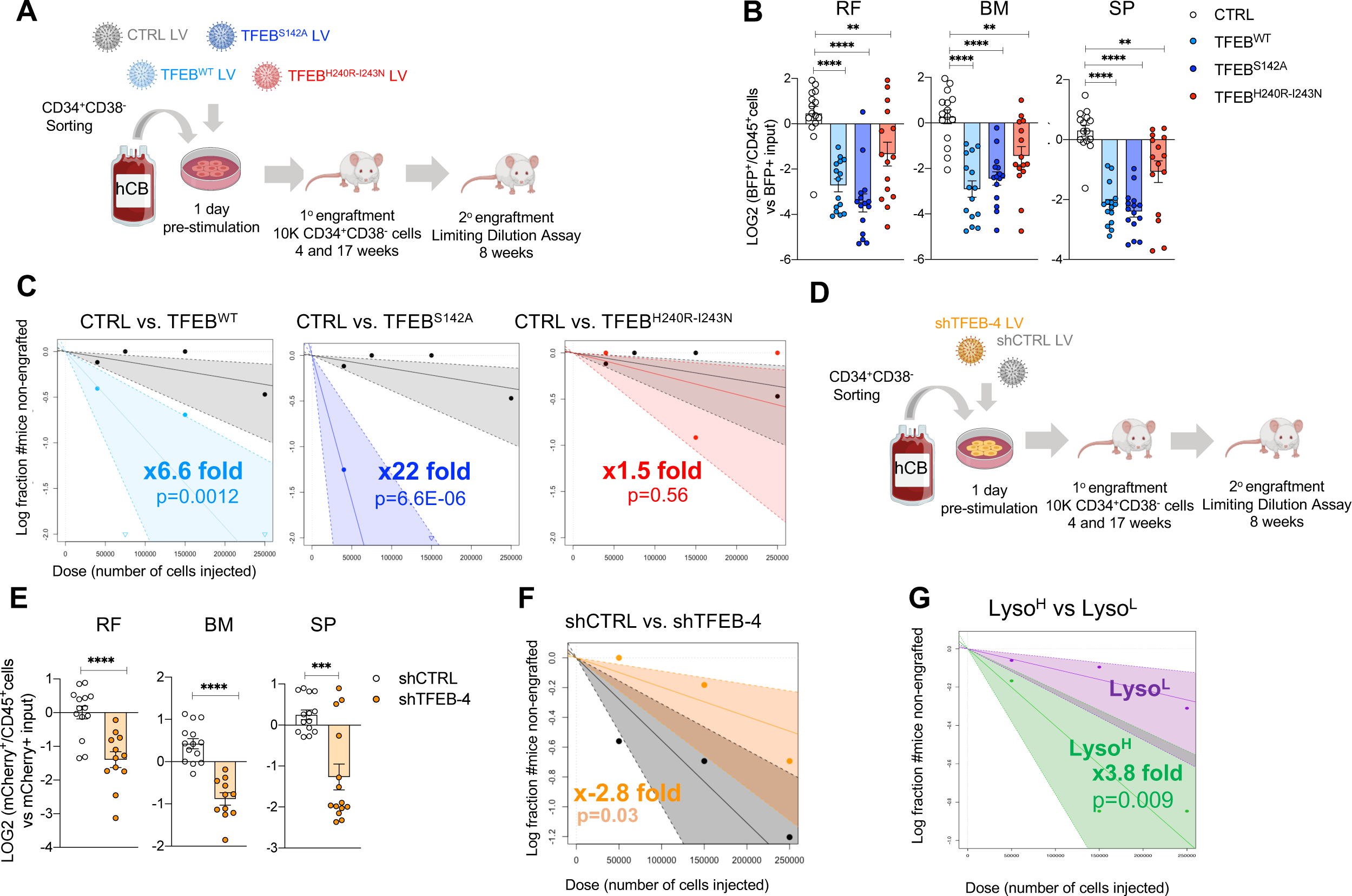
TFEB-directed lysosomal activity controls LT-HSC self-renewal. **A.** Experimental scheme: CD34^+^CD38^-^ cells isolated from hCB were transduced with lentiviral vectors expressing BFP and one of the following genes: CTRL, TFEB^WT^, TFEB^S142A^ or TFEB^H240R-I243N^. At 1d post-transduction cells were intrafemorally injected into 8-10w old NSG- W41 and NSG mice. Male NSG mice were sacrificed at 4w, female NSG and male NSG-W41 mice at 17w post-xenotransplantation for analysis. CD45^+^/BFP^+^ cells were sorted from primary NSG mice at 17w for secondary limiting dilution assays (LDA) in NSG-GM3 mice. NSG-GM3 mice were sacrificed at 8w post-xenotransplantation for stem cell frequency analysis. **B.** From (A): LOG2 ratio of %BFP^+^ in hCD45^+^ cells at 17w of engraftment in NSG-W41 mice vs input (3d post-transduction). **C.** Stem cell frequency calculated from LDA. See Tables in Fig. S10L,M**. D.** As in (A) using lentivirus expressing mCherry and shCTRL or shTFEB. **E.** From (D): LOG2 ratio of %mCherry^+^ in hCD45^+^ cells at 17w of engraftment in NSG-W41 mice vs input (3d post- transduction). **F.** Stem cell frequency calculated from LDA. See Tables in Fig. S11F,G**. G.** Stem cell frequency calculated from LDA. See Tables in Fig. S12D,E. Each *in vivo* experiment was performed with three independent CB per experiment with 4-5 mice/CB. (RF: right femur, injected bone; BM: left femur and right and left tibiae; SP: spleen). *P<0.05, **P<0.01, ***P<0.001. Unpaired t-test unless otherwise indicated.

Next, we examined the effects of modulating TFEB activity through altering the phosphorylation of the S142 and S211 sites that are known to be targeted by several kinases (mTORC1, CDK4/6, ERK1/2)(*19, 31*). In nutrient-rich conditions, S142 and S211 phosphorylation of TFEB results in its cytoplasmic retention, whereas upon nutrient starvation, dephosphorylated TFEB translocates to the nucleus to induce the expression of lysosomal genes(*13, 19, 31*). Overexpression of a constitutively nuclear form of TFEB (TFEB^S142A^), generated through disruption of the S142 phosphorylation site, in CD34^+^CD38^-^ hCB cells led to significant reduction of graft size in primary transplanted mice but a large 22-fold increase in the frequency of self-renewing stem cells (**Fig. 5A-C;Fig. S9A-F;Fig. S10G-M)**. The greater enrichment of stem cell frequency in TFEB^S142^ compared to TFEB^WT^-OE cells suggests that regulation of TFEB subcellular localization is important for LT-HSC self-renewal (**Fig. 5A- C;Fig. S10G-M**).

Silencing TFEB in CD34^+^CD38^-^ hCB cells also resulted in smaller graft size in primary mouse recipients, but in contrast to the findings with TFEB-OE vectors, the reduced engraftment in the shTFEB group was now accompanied by a 2.8-fold reduction in stem cell frequency by serial LDA (**Fig. 5D-F;Fig. S11A-G)**. Importantly, the myeloid and erythroid bias induced by TFEB-OE and TFEB-knockdown, respectively, was recapitulated in xenotransplantation experiments (**Fig. S11H-J**). Interestingly, Lyso^H^ LT-HSC exhibited a 3.8-fold increase in stem cell frequency despite showing a small reduction in primary graft size compared to Lyso^L^ LT- HSC, strengthening the link between lysosomal activity and LT-HSC stemness potential (**Fig. 5G; Fig. S12A-E**). In summary, LT-HSC self-renewal requires the TFEB-regulated lysosomal program to limit LT-HSC metabolic activity and expansion, whereas inhibition of this program leads to LT-HSC activation followed by exhaustion.

## Discussion

Here, we have identified an organelle-based model of stem cell fate determination where TFEB and MYC balance the activity of lysosomes to regulate the self-renewal and differentiation properties of human LT-HSC. In an unperturbed homeostatic setting, TFEB (normally implicated in stress responses) induces constitutive lysosomal flux in LT-HSC that actively maintains quiescence, preserves self-renewal and governs lineage commitment. These effects are tied to endolysosomal degradation of membrane receptors, pointing to a role for TFEB in coordinating how LT-HSC sense environmental changes and initiate the earliest steps of their fate transitions and lineage commitment decisions. Such transitions are delicately balanced by a TFEB/MYC dichotomy where MYC is a driver of LT-HSC anabolism and activation, counteracting TFEB function by serving as a negative transcriptional regulator of lysosomes; conversely TFEB counteracts MYC activity (**Fig. S12F**). TFEB activation is post-transcriptionally regulated through phosphorylation and subcellular localization by upstream kinases and results in the limiting of LT-HSC quiescence and self-renewal.

Our findings point to a mechanism whereby the TFEB-induced lysosomal program actively maintains LT-HSC in a quiescent state via endosomal trafficking of external sensing machinery including signaling and nutrient-uptake receptors to lysosomes for degradation, as exemplified here by the iron transporter TfR1. We hypothesize that this endolysosomal system renders quiescent LT-HSC more refractory to specific environmental cues in order to restrain aberrant mitogenic and anabolic activation and prevent LT-HSC exhaustion, as shown by our TFEB and TfR1 knockdown experiments. Quiescent mouse HSC express lower membrane levels of TfR1 compared to progenitor cells, which require higher levels of iron-uptake for proliferation and its deletion resulted in profound reconstitution defects(*32*). Regulation of TfR1 by lysosomes in qLT-HSC may have a protective role by preventing iron overload and the resultant ROS production that would be detrimental to HSC self-renewal(*33*). Our findings on TfR1 also help explain an emerging body of work on how lineage determination often occurs already within the stem cell compartment(*21*). Active suppression of TFEB and its downstream lysosomal degradation of TfR1 within LT-HSC is required for commitment along the erythroid lineage: indeed, activation of TFEB can abolish erythroid differentiation even after lineage commitment has occurred. Our transcriptomic analysis of TFEB-OE LT-HSC indicates that endolysosomal degradation of other membrane receptors such as IGF1R (insulin receptor), which is upstream of the PI3K-Akt-mTORC1 pathway, could also be taking place in qLT-HSC, a finding consistent with observed EGFR degradation in neural stem cells(*34*). Thus, it is highly likely that the dichotomous TFEB/MYC-mediated control of lysosomal activity will be relevant in other tissue-specific adult stem cells such as neural and muscle stem cells that are maintained in quiescence for prolonged periods of time. In summary, our work sheds light on the transcriptional control of lysosomes and its crucial r ole in stemness regulation, opening new routes to explore in regenerative medicine.

## Acknowledgments

We thank the obstetrics units of Trillium Health, William Osler and Credit Valley Hospitals for CB; The UHN-Sickkids Flow cytometry facility for cell sorting; The Advance Optical Microscope Facility (AOMF) for confocal microscopy, Linda Penn, Jason DeMelo and Aaliya Tamachi for the MYC overexpressing vector and all lab members of the Dick lab for critical feedback.

## Author contributions

L.G.P conceived the study, performed research, analyzed data, and wrote the manuscript; F.S and W.W performed research and analyzed data; J.C, O.G, J.M, M.S, K.P, G.K, K.G, S.T and E.W performed research; V.V, A.M, M.C.S.Y and A.G.X.Z analyzed data; K.B.K and S.Z.X performed research and wrote the manuscript; and J.E.D wrote the manuscript, secured funding and supervised the study.

## Material and Methods

### Methods Table

| REAGENT or RESOURCE | SOURCE | IDENTIFIER |
| --- | --- | --- |
| **Antibodies** |  |  |
| FITC–anti-CD45RA | BD | 555488 |
| PE–anti-CD90 | BD | 555596 |
| PECy5–anti-CD49f | BD | 551129 |
| V450–anti-CD7 | BD | 642916 |
| PECy7–anti-CD38 | BD | 335790 |
| APC–anti-CD10 | BD | 340923 |
| APCCy7–anti-CD34 | BD | custom made by<br>BD |
| BV711-CD19 | BD | 563036 |
| Alexafluor700–anti-CD7 | BD | 561603 |
| Alexafluor700–anti-CD10 | BD | 563500 |
| BV510-CD45 | BD | 563891 |
| biotin-Flt3L | BD | N/A |
| Streptavidin-Qdot605 | ThermoFisher | Q10101MP |
| Streptavidin-PE | BD | 554061 |
| V450-anti-CD15 | BD | 642917 |
| FITC-anti-CD3 | 349201 | 349201 |
| V500-anti-CD45 | BD | 560777 |
| PE–anti-CD19 | BD | 349209 |
| PE–anti-GlyA | Beckman Coulter | A07792 |
| BV786–anti-CD33 | BD | 740974 |
| PECy5–anti-CD45 | Beckman Coulter | A07785 |
| PECy7–anti-CD14 | Beckman Coulter | A22331 |
| APC–anti-CD33 | BD | 551378 |
| APC–anti-CD45 | BD | 340943 |
| FITC–anti-CD45 | BD | 347463 |
| V421–anti-CD10 | BD | 562902 |
| BV605 anti-CD56 | BD | 562780 |
| PE-anti-CD71 | BD | 555537 |
| APCCy7 anti-CD41 | Beckman Coulter | 6607115 |
| BV786-anti-CD71 | BD | 563768 |
| Ki67-PE | BD | 556027 |
| Anti-TFEB | R&D biosystems | MAB9170-100 |
| Anti- MYC Alexa647 | R&D biosystems | IC36961R-100ug |
| Anti-LAMP1 Alexa488 | R&D biosystems | IC7985G |
| Anti-CD71 APC | BD | 551374 |
| Purified Mouse Anti-Human CD71 | BD | 555534 |
| Human Cathepsin D Antibody | R&D biosystems | AF1014 |
| Goat Anti-mouse secondary antibody Alexa568 | Thermo Fisher | A-11004 |
| Goat Anti-rabbit secondary antibody Alexa488 | Thermo Fisher | A-11008 |
| **Biological Samples** |  |  |
| Human umbilical cord blood samples | Trillium, Credit Valley and Brampton Civic Hospital | N/A |
| **Chemicals, Peptides, and Recombinant Proteins** |  |  |
| DNase I | Roche | 11284932001 |
| Ammonium Chloride | Stem Cell Technologies | 07850 |
| FLT3 Ligand | Milteny Biotec | 130-096-474 |
| IL6 | Milteny Biotec | 130-093-934 |
| SCF | Milteny Biotec | 130-096-696 |
| TPO | Milteny Biotec | 130-095-752 |
| EPO | Janssen | Eprex 10,000 IU/ml |
| IL3 | Milteny Biotec | 130-095-068 |
| GM-CSF | Milteny Biotec | 130-093-866 |
| AmpliTaq Gold 360 Polymerase | ThermoFisher | 4398813 |
| BamH1-HF | NEB | R0136S |
| Mlu1-HF | Invitrogene | 15432-016 |
| KOD Hot Start DNA Polymerase | Millipore-Sigma | 71842 |
| DpnI | New England Biolabs | R0176S |
| Paraformaldehyde 16% Solution (methanol-free) | Agar scientific | AGR1026 |
| Poly-L-lysine solution | Sigma | P8920 |
| IGEPAL® CA-630 | Sigma | I8896-50ML |
| DMSO | Fisher Chemical | D128-500 |
| Bafilomycin A1 | Sigma | B1793 |
| MYC inhibitor | Milipore-sigma | 475956-10MG |
| DAPI | Sigma | 10236276001 |
| Fluoromount G | Thermo Fisher | 00-4958-02 |
| Tween 20 | Sigma | P9416-100ML |
| Triton X-100 | Sigma | T8787-100 |
| **Commercial Assays** |  |  |
| StemSep™ Human Hematopoietic Progenitor Cell Enrichment Kit | Stem Cell Technologies | 14066 |
| CD34 MicroBead Kit, human | Milteny Biotec | 130-097-047 |
| LS Columns | Milteny Biotec | 130-042-401 |
| Mouse Cell Depletion Kit | Milteny Biotec | 130-104-694 |
| SuperScript VILO | ThermoFisher | 1754050 |
| Power SYBR Green | ThermoFisher | 4367659 |
| Magic Red™ Cathepsin B Kit | BioRad | ICT937 |
| RNeasy Micro Kit | Qiagen | 74004 |
| PicoPure® RNA Isolation Kit | Thermo Fisher | KIT0204 |
| Gateway™ LR Clonase™ II Enzyme Mix | Thermo Fisher | 11791020 |
| CYTO-ID® Autophagy detection kit | Enzo Life Sciences | ENZ-51031-K200 |
| RNase-Free DNase Set (50) | Qiagen | 79254 |
| Click-iT® Plus EdU Alexa Fluor® 647 Flow Cytometry | Thermo Fisher | C10634 |
| CellROX Deep Red | ThermoFisher | C10422 |
| CellROX Orange | ThermoFisher | C10443 |
| MitoTracker Green | ThermoFisher | M7514 |
| LysoTracker Blue | ThermoFisher | L7525 |
| LysoTracker Green | ThermoFisher | L7526 |
| LysoTracker Deep Red | ThermoFisher | L12492 |
| LysoSensor Green DND-189 | ThermoFisher | L7535 |
| RNeasy Plus Micro Kit | Qiagen | 74034 |
| Sytoxblue | ThermoFisher | S11348 |
| Propidium Iodide | ThermoFisher | P3566 |
| BD cytofix/cytoperm fixation solution | BD | 554722 |
| BD PermWash | BD | 554723 |
| Phusion High fidelity PCR master Mix with HF buffer | NEB | M0531S |
| MiniElute PCR purification Kit | Qiagen | 28004 |
| Nextera DNA library prep kit 96 samples | Illumina | 15028211 |
| Syber Green | ThermoFisher | S7563 |
| Hoechst 33342 | BD | 561908 |
| **Other** |  |  |
| X-vivo medium | Lonza | 04-380Q |
| H5100 media | StemCell Technologies | 05150 |
| Fetal bovine serum | Sigma | F1051-500mL |
| Iscove's modified Dulbecco's medium (IMDM) | Gibco | 12440-053 |
| MethoCult™ Optimum | Stem Cell Technologies | H4034 |
| StemPro™-34 SFM | Gibco | 10640-019 |
| StemPro nutrients | Gibco | 10641-025 |
| L-glutamine | Multicell | 609-065-EL |
| Pen/Strep | Gibco | 15140-122 |
| Human LDL | Stem Cell Technologies | 02698 |
| 8 Well Chamber, removable | ibidi | 80841 |
| 12 Well Chamber, removable | ibidi |  |

### Lead contact and materials availability

Further information and requests for resources and unique/stable reagents generated in this study should be directed to and are available from the Lead Contact, John E. Dick, but a completed Materials Transfer Agreement may be required.

## Method details

### Human cord blood samples

Human CB samples were obtained with informed consent from Trillium Health, Credit Valley and William Osler Hospitals according to procedures approved by the University Health Network (UHN) Research Ethics Board. Mononuclear cells (MNC) from pools of male and female CB units (∼4-15 units) were obtained by centrifugation on Lymphoprep medium, and after ammonium chloride lysis MNC were enriched for CD34^+^ cells by positive selection with the CD34 Microbead kit and LS column purification with MACS magnet technology (Miltenyi). Resulting CD34^+^ CB cells were viably stored in 50% PBS, 40% fetal bovine serum (FBS) and 10% DMSO at -80°C or −150 °C.

### Mice

Animal experiments were done in accordance with institutional guidelines approved by University Health Network Animal care committee. All in vivo experiments were done with 8- to 12-week-old female/male *NOD.Cg-PrkdcscidIl2rgtm1Wjl/SzJ* (NSG) mice (JAX) and 8- to 12-week-old female/male *NOD.Cg-PrkdcscidIl2rgtm1WjlTg(CMV-IL3,CSF2,KITLG)1Eav/MloySzJ* (NSG-SGM3) that were sublethally irradiated with 225 cGy, 24 h before transplantation, or with 8- to 12-week-old male *NOD.Cg- PrkdcscidIl2rgtm1WjlKitem1Mvw/SzJ* (NSGW41) mice that were not irradiated. All mice were housed at the animal facility (ARC) at Princess Margaret Cancer Centre in a room designated only for immunocompromised mice with individually ventilated racks equipped with complete sterile micro-isolator caging (IVC), on corn-cob bedding and supplied with environmental enrichment in the form of a red house/tube and a cotton nestlet. Cages are changed every <7 days under a biological safety cabinet. Health status is monitored using a combination of soiled bedding sentinels and environmental monitoring.

### Flow Cytometric Analysis and Sorting

#### Sorting

CD34^+^ and CD34^-^ human CB cells were thawed via slow dropwise addition of X-VIVO 10 medium with 50% FBS and DNaseI (200 μ /ml). Cells were spun at 350g for 10min at 4 °C g and then resuspended in PBS+2.5% FBS. For all in vitro and in vivo experiments, the full stem and progenitor hierarchy sort was performed as described in Notta et al.(*21*) and shown in Fig. S1A and Fig. S7B. Cells were resuspended in 100 lμ per 1x10^6^ cells and stained in two subsequent rounds for 15 min at room temperature each. See Table 1 for antibodies.

#### Cell cycle

Cells were cultured with EdU for 1 hour and harvested for EdU Click-it reaction following manufacturer’s instructions. Cells were then incubated with PE-anti-Ki67 antibody in PermWash solution overnight at 4°C. Prior to flow cytometry analysis cells were stained with Hoechst 33342.

#### Cell dyes

Cells were incubated for 30 min at 37**°**C with the indicated dyes listed in Table 1 following manufacturer’s instructions (CellROX, 5μ; MitoTracker, 0.1μ; LysoTracker, 75nM; Magic Red, 1/260 dilution; CytoID, 1/500 dilution; LysoSensor, 1/500 dilution) in respective culture media. After staining, cells were washed once and resuspended in PBS+2.5% FBS and analyzed by flow cytometry.

BD sorters FACSAria II, FACSAria III and FACSAria FUSION; and BD analyzers FACSCelesta were used.

### Immunostaining analysis

Cells were spun onto Poly-L-Lysine-coated slides (200 xg, 10 min), fixed with 4% paraformaldehyde and permeabilized with 0.5% Triton before blocking (PBS, 10% FBS, 5% BSA). Slides were incubated with primary antibodies listed in Table 1 in blocking solution O/N at 4°C. Secondary antibodies also listed in Table 1 were added in PBS, 0.025% Tween for 1.5 h at room temperature (RT). After washing, nuclei were stained with 1 μ/mL DAPI and slides g were mounted (Fluoromount G). Single cell images were captured by a Zeiss LSM700 Confocal Microscope (oil, 63x/1.4NA, Zen 2012) and processed and analyzed with ImageJ/Fiji and FlowJo10. Antibody dilutions used are the following: LAMP-1 1/200; TFEB 1/50; MYC 1/100; TfR1 1/50; Cathepsin D 1/100.

#### Analysis of TFEB-MYC correlation in single cell immunostaining data

fluorescence intensities for LAMP1, nuclear DAPI, MYC, nuclear MYC, TFEB, and nuclear TFEB were normalized to 10,000 for each cell and subject to log transformation.

### Time-lapse imaging

Time-lapse experiments were conducted at 37°C, 5% O_2_, 5%CO_2_ on μ-slide VI^0.4^ channel slides (IBIDI) coated with 20 μg/ml anti-human CD43-biotin antibody(*37*). BFP^+^ cells were sorted 3 days post-lentiviral transduction and cultured overnight in phenol red free IMDM supplemented with 20% BIT (Stemcell Technologies), 100 ng/mL human recombinant Stem Cell Factor (SCF), 50 ng/mL human recombinant Thrombopoietin (TPO), 100 ng/mL human Fms-related tyrosine kinase 3 ligand (Flt3L, all R&D Systems), 2 mM L-GlutaMAX (Gibco), 100 U/mL penicillin and 100 µg/mL Streptomycin (Gibco) before imaging. Brightfield images were acquired every 8 minutes for 3-4 days using a Nikon-Ti Eclipse equipped with a Hamamatsu Orca Flash 4.0 camera and a 10x CFI Plan Apochromat λ objective (NA 0.45). Single-cell tracking and fate assignment were performed using self-written software as previously described(*11, 35, 36*). Time to division was calculated using R 3.5.3.

### Cloning of lentiviral overexpression constructs

Lentiviral constructs expressing *TFEB* variants were constructed by Gateway^TM^ cloning from pENTR223 donor plasmids into a lentiviral pRRL-based and Gateway^TM^ adapted vector (pLBC2-BS-RFCA(*37*)) downstream of a SFFV promoter and upstream of tagBFP driven by an EFS/SV40 chimeric promoter using the Gateway^TM^ LR Clonase^TM^ II Enzyme Mix according to manufacturer’s instructions (Thermo Fisher). The original donor plasmid pENTR223-TFEB (HsCD00373101) was obtained from PlasmID (DF/HCC DNA Resource Core at Harvard Medical School). By standard site-directed PCR mutagenesis a STOP-codon was inserted (fw primer: 5’-TAG ACC CAG CTT TCT TGT ACA AAG TTG-3’; rev primer: 5’-TCA CAG CAC ATC GCC CTC C-3’) and subsequently pENTR223-TFEB-S142A (fw primer: 5’- CAC CCA TGG CCA TGC TGC AC-3’; rev primer: 5’- CAT TGG GAG CAC TGT TGC CAG C-3’) and pENTR223-TFEB- H240R_ I243N (fw primer: 5’- CGG AAC TTA AAT GAA AGG AGA CGA AGG; rev primer: 5’- ATT GTC TTT CTT CTG CCG CTC C-3’) were generated using KOD Hot Start DNA Polymerase (Millipore-Sigma) followed by DpnI (NEB) digestion. The negative control (CTRL) vector for overexpression encodes *gp91phoxP415H* (catalytic inactive gp91phox/*CYBB*; (*35*) that was previously cloned into the same lentiviral Gateway^TM^ adapted vector (pLBC2-BS-RFCA(*37*)). MYC overexpressing vector was kindly provided by Linda Penn’s lab.

### Cloning of lentiviral knockdown constructs

The negative control (CTRL) vector for knockdown is a shRNA directed against Renilla luciferase (shCTRL)(*37*). Additional shRNA sequences were predicted using the Sherwood algorithm as in(*38*) and ordered as Ultramer DNA oligos (IDT). Subsequently, shRNAs were amplified using AmpliTaq Gold 360 Polymerase (ThermoFisher) using shRNA amplification Forward and Reverse primers. The PCR product was subsequently digested with BamH1 and Mlu1 and cloned into a UltramiR scaffold (miR30) within a pRRL-based vector downstream of a SFFV promoter and upstream stream of mCherry (pLBC2-mCherry).

#### shRenilla

TGCTGTTGACAGTGAGCGCAGGAATTATAATGCTTATCTATAGTGAAGCCACAGATGTATAGATAAGCATTATAATTCCTATGCCTACTGCCTCGGA1

#### shTFEB-2

TGCTGTTGACAGTGAGCGAACGATGTCCTTGGCTACATCATAGTGAAGCCACAGATGTATGATGTAGCCAAGGACATCGTCTGCCTACTGCCTCGGA

#### shTFEB-3

TGCTGTTGACAGTGAGCGAGATGATGTCATTGACAACATATAGTGAAGCCACAGATGTATATGTTGTCAATGACATCATCCTGCCTACTGCCTCGGA

#### shTFEB-4

TGCTGTTGACAGTGAGCGCAGAAAGACAATCACAACTTAATAGTGAAGCCACAGATGTATTAAGTTGTGATTGTCTTTCTTTGCCTACTGCCTCGGA

#### shLAMP-1

TGCTGTTGACAGTGAGCGATGGACGAGAACAGCATGCTGATAGTGAAGCCACAGATGTATCAGCATGCTGTTCTCGTCCAGTGCCTACTGCCTCGGA

#### shLAMP-2

TGCTGTTGACAGTGAGCGCCACGAGAAATGCAACACGTTATAGTGAAGCCACAGATGTATAACGTGTTGCATTTCTCGTGATGCCTACTGCCTCGGA

#### shLAMP-3

TGCTGTTGACAGTGAGCGACAGCTCATGAGTTTTGTTTAATAGTGAAGCCACAGATGTATTAAACAAAACTCATGAGCTGGTGCCTACTGCCTCGGA

#### shLAMP-4

TGCTGTTGACAGTGAGCGCGACTGTGGAATCTATAACTGATAGTGAAGCCACAGATGTATCAGTTATAGATTCCACAGTCTTGCCTACTGCCTCGGA

#### shTFRC-1

TGCTGTTGACAGTGAGCGCGAACCAGATCACTATGTTGTATAGTGAAGCCACAGATGTATACAACATAGTGATCTGGTTCTTGCCTACTGCCTCGGA

#### shTFRC-2

TGCTGTTGACAGTGAGCGCCCAGATCACTATGTTGTAGTATAGTGAAGCCACAGATGTATACTACAACATAGTGATCTGGTTGCCTACTGCCTCGGA

#### shTFRC-3

TGCTGTTGACAGTGAGCGCCTCAATGATCGTGTCATGAGATAGTGAAGCCACAGATGTATCTCATGACACGATCATTGAGTTGCCTACTGCCTCGGA

#### shTFRC-4

TGCTGTTGACAGTGAGCGCCAGCCAACTGCTTTCATTTGATAGTGAAGCCACAGATGTATCAAATGAAAGCAGTTGGCTGTTGCCTACTGCCTCGGA

### Lentiviral Production and Transduction

VSV-G pseudotyped lentiviral vector particles were produced and titers were determined as previously described(*37*). Unless stated otherwise, after 16-20 hours of pre-stimulation in low cytokine media (see descriptioon below) cells were transduced with lentiviral vectors described above at matching multiplicity of infection(*37*) aiming at mid-range (20-40%) transduction efficiencies but without lentiviral preparation exceeding 20% of total culture volume. Transduction efficiency (%BFP^+^ or %mCherry^+^) was determined at day 3 post-transduction by flow cytometry on a BD Celesta, which served as initial input estimate for xenotransplantation assays.

### Cell Culture

For *in vitro* experiments sorted cells were cultured in 96 well-plate round bottom with the indicated cell media.

- <u>high cytokine media</u>: StemPro-34 SFM media with the supplied supplement, 1x L-Glutamine, 1x Pen/Strep, 0.02% Human LDL and cytokines FLT3L (20 ng/mL), GMCSF (20 ng/mL), SCF (100 ng/mL), TPO (100 ng/mL), EPO (3 U/mL), IL-3 (10 ng/mL), IL-6 (50 ng/mL). This media was used to culture cells with the presence of DMSO, MYC inhibitor or BAF.

- <u>low cytokine media</u>: X-Vivo 10, 1% BSA, L-Glutamine, Pen/Strep and cytokines SCF (100 ng/ml), Flt3L (100 ng/ml), TPO (50 ng/ml) and IL7 (IL-7; 10 ng/ml). This media was used to transduce cells as indicated above.

For drug studies of untransduced cells, DMSO or MYC inhibitor (53.3 μM) were added at the time of seeding in high cytokine media and cells were harvested 4 days later for indicated analysis.

For drug studies of transduced cells, DMSO, MYC inhibitor (53.3 μM) or Bafilomycin (20 nM) were added in high cytokine media 1day after transduction with lentiviral vectors. Cells were harvested 4 days later for indicated analysis.

For lysosomal turnover studies, Bafilomycin (20 nM) or equivalent amount of DMSO was added to the cultures for 3 hours in StemPro-34 SFM media (1x L-Glutamine, 1x Pen/Strep, 0.02%) without cytokines.

### Colony-Forming Cell Assays

#### For Lyso^H^ and Lyso^L^ cells

∼ 20K LT-HSC were cultured for 4 days in high cytokine media and stained with LysoTracker as described above. For each group (based on LysoTracker fluorescence) 150 cells were sorted directly into 2 mL methylcellulose, supplemented with FLT3 Ligand (10 ng/mL) and IL6 (10 ng/mL) and plated onto 2x35 mm dishes as duplicates.

#### For transduced cells

At day 3 post-transduction, 150 LT-HSC BFP^+^ or mCherry^+^, 200 ST-HSC BFP^+^, 300 MEP F1 BFP^+^ or 300 MEP F2/3 BFP^+^ cells were sorted directly into 2 mL methylcellulose, supplemented with FLT3 Ligand (10 ng/mL) and IL6 (10 ng/mL) and plated onto 2x35 mm dishes as duplicates.

Colonies were allowed to differentiate for 10-11 days and then morphologically assessed in a blind fashion by a second investigator. Subsequently, colonies from replicate plates were pooled, resuspended in PBS/5% FBS and stained for flow cytometry analysis. For serial CFC assays, 0.5% of progeny from LT-HSC was added to fresh methylcellulose as above, replated and scored after 10-11 days.

### Single-Cell Stromal Assays

Single cell in vitro assays were set up as described previously(*39*) with low passage murine MS- 5 stroma cells seeded at a density of 1500 cells per 96-well and grown for 2-4 days in H5100 media. One-day prior to coculture initiation, the H5100 media was removed and replaced with 100 μl erythro-myeloid differentiation media: StemPro-34 SFM media with the supplied supplement, 1x L-Glutamine, 1x Pen/Strep, 0.02% Human LDL and the following cytokines: FLT3L (20 ng/mL), GMCSF (20 ng/mL), SCF (100 ng/mL), TPO (100 ng/mL), EPO (3 U/mL), IL-2 (10 ng/mL), IL-3 (10 ng/mL), IL-6 (50 ng/mL), IL-7 (20 ng/mL), and IL-11 (50 ng/mL). Sorted single-cells were deposited in each well (80 wells/96-well plate). Colonies were scored after 15-17 days under the microscope and every individual well containing a visible colony was stained with antibodies listed in Table1 and analyzed by flow cytometry using a BD FACSCelesta instrument equipped with a high throughput sampler (HTS).

### Xenotransplantation

The progeny of 10K CD34^+^CD38^-^ transduced cells one day after transduction were intra-femoral injected in aged and gender matched 8-12 wk old male and female recipient NSG, NSG-SGM3 or NSG-W41 mice. At indicated time points, mice were euthanized, injected femur and other long bones (non-injected femur, tibiae) were flushed separately in Iscove’s modified Dulbecco’s medium (IMDM)+5%FBS and 5% of cells were analyzed for human chimerism along with BFP or mCherry and antibodies listed in Table 1. Sick and miss-injected mice were excluded from analysis. For purification of human cells from xenotransplanted mice, fresh or thawed BM from individual or from pools of 2-5 mice were mouse depleted (Mouse Cell Depletion Kit, Miltenyi) according to manufacturer’s protocol and CD45^+^/BFP^+^ or CD45^+^/mCherry^+^ were sorted for serial transplantation by intra-femur injection in NSG-SGM3 mice at the indicated cell doses. A mouse was considered engrafted if % of human CD45^+^ cells > 0.1. For LDA experiments LT-HSC frequency was estimated using the ELDA software (http://bioinf.wehi.edu.au/software/elda/).

For <u>Lyso^H^ and Lyso^L^ LT-HSC:</u> ∼ 20K LT-HSC were cultured for 4 days in high cytokine media and stained with LysoTracker as described above. For each group (based on LysoTracker fluorescence) 1K LT-HSC were intra-femoral injected in aged and gender matched 8-12 wk old male and female recipient NSG-W41 or NSG-SGM3 mice and. Engraftment analysis of primary and secondary xenograft was performed as described above.

### RNA-seq processing and analysis

Freshly sorted populations from 3-5 independent CB pools or transduced cells in vitro after a second sort (PI^-^BFP^+^ or PI^-^mCherry^+^) on day 3 or 6 post-transduction were directly resuspended and frozen (-80°C) in PicoPure RNA Isolation Kit Extraction Buffer. RNA was isolated using the PicoPure RNA Isolation Kit (Thermo Fisher) according to manufacturer’s instructions. Samples that passed quality control according to integrity (RIN>8) and concentration as verified on a Bioanalyzer pico chip (Agilent Technologies) were subjected to further processing by the Center for Applied Genomics, Sick Kids Hospital: cDNA conversion was performed using SMART-Seq v4 Ultra Low Input RNA Kit for Sequencing (Takara) and libraries were prepared using Nextera XT DNA Library Preparation Kit (Illumina). Equimolar quantities of libraries were pooled and sequenced with 4 cDNA libraries per lane on a High Throughput Run Mode Flowcell with v4 sequencing chemistry on the Illumina 30 HiSeq 2500 following manufacturer’s protocol generating paired-end reads of 125-bp in length to reach depth of 55-75 million reads per sample. Reads were then aligned with STAR v2.5.2b against hg38 and annotated with ensembl v90. Default parameters were used except for the following: chimSegmentMin 12; chimJunctionOverhangMin 12; alignSJDBoverhangMin 10; alignMatesGapMax 100000; alignIntronMax 100000; chimSegmentReadGapMax parameter 3; alignSJstitchMismatchNmax 5 -1 5 5. Read counts were generated using HTSeq v0.7.2 and general statistics were obtained from picard v2.6.0. Differential gene expression was performed using edgeR_3.24.3 following recommended practices. For the qLT-HSC vs aLT-HSC comparison (Supplementary Table S1) the LT-HSC from the dataset GSE125345 were compared to the cultured CTRL LT-HSC from the TFEB-OE experiment. For the aLT-HSC vs aST-HSC comparison (Supplementary Table S8), CTRL LT-HSC and ST-HSC from the TFEB-OE experiment were compared. These data were used in a single sample gene set variation analysis using TFEB-OE and MYC-OE up and downregulated genes (top 100, 250 and 500 genes) using the gsva function of the R package GSVA_1.30.0.

Conditional Quantile Normalization (CQN) from the cqn R package was applied on TMM normalized raw counts in order to correct for gene length effect. Expression of TFEB, TFEC, TFE3 and MITF was extracted from the data and plotted in each population using R stripchart.

### Transcriptomic analysis of published datasets

Several transcriptomics data containing HSC and other hematopoietic populations from cord blood (CB), bone marrow (BM), mobilized peripheral blood cells (mPB) or fetal liver (FL) tissues were obtained from Gene Expression Omnibus data portal (GEO).

#### GSE42414 (Laurenti et al. 2013, PMID: 23708252)(*30*)

Illumina HumanHT-12 beadchip expression data from lineage-depleted CB cells were downloaded from GEO. Normalized log2-transformed signal was used in a moderated t test in the R limma_3.38.3 package to compare HSC1 (Lin- CD34+ CD38- CD45RA- CD90+ CD49f+) and MPP Lin- CD34+ CD38- CD45RA- CD90- CD49f-). A rank file was obtained by ordering all genes using the t statistics from upregulated to downregulated genes in HSC1 compared to MPP and GSEA_4.0.3 was run using default parameters using the KEGG lysosome as gene-set.

#### GSE76234 (Notta et al. 2016, PMID: 26541609)(*21*)

FPKM expression data from Illumina HiSeq 2000 RNA sequencing aligned on hg19 were downloaded from GEO. Median gene expression for HSC, MPP(+-), MPP(++), and MPP(--) lineage-depleted CB populations were available and used for this analysis. These data were used in a single sample gene set variation analysis with the KEGG-Lysosome gene-set using the gsva function of the R package GSVA_1.30.0. They were also used to plot the expression of TFEB, TFEC, TFE3 and MITF in each population.

#### GSE109093 (Cesana et al. 2018, PMID: 29625070)(*20*)

FPKM expression data from Illumina HiSeq 2000 RNA sequencing aligned on hg19 were downloaded from GEO. Median gene expression for HSC (CD34+ CD38– CD90+ CD45RA–) and committed CD34+CD38+ progenitor populations (PROG) from BM, FL and CB were available and used for this analysis. These data were used to plot the expression of TFEB, TFEC, TFE3 and MITF in each population.

#### GSE17054, GSE19599, GSE11864, E-MEXP-1242 (Hemaexplorer)

Log2 batch corrected expression signal from Affymetrix Human Genome U133 Plus 2.0 Array were downloaded from Bloodspot (http://servers.binf.ku.dk/bloodspot/). It includes normal hematopoietic populations from BM and peripheral blood. Expression of TFEB, TFEC, TFE3 and MITF was extracted from the data and plot in each population using R stripchart.

### Pathway enrichment analysis and visualization

Pathway enrichment analysis and visualization was performed as described previously(*40*). Briefly, a score to rank genes from top up-regulated to down-regulated was calculated using the formula -sign(logFC) * -log10(pvalue). The rank file from each comparison was used in GSEA analysis (http://software.broadinstitute.org/gsea/index.jsp) using 2000 permutations and default parameters against indicated gene sets. All gene sets were obtained from a pathway database http://baderlab.org/GeneSets. EnrichmentMap version 3.1.0 in Cytoscape 3.7.0 was used to visualize enriched gene-sets with indicated FDR-q value and NES and a Jaccard coefficient set to 0.375. Full results in the comparison of CTRL vs TFEBWT-OE LT-HSC can be seen in Supplementary Table 3. g:profiler (https://biit.cs.ut.ee/gprofiler/gost) was performed using the upregulated genes by TFEB-OE (FDRq-value<=0.01), results can be seen in Supplementary Table 3.

### scRNA-seq Signature Enrichment

Using signatures derived from over-expression of TFEB and MYC in LT-HSCs (FDR < 0.05), we scored single CD34+CD38- HSPCs from three publicly available datasets for their relative expression of each signature. We then compared signature enrichment between cell cycle-primed HSPCs and non-primed HSPCs, as defined in Xie et al.(*7*) The MYC OE signature was higher in the cycle-primed HSPCs while the TFEB OE signature was higher in the non-primed HSPCs, and these results were consistent across all three datasets(*26,27,28*).

### Comparison of TFEB-regulated genes in MEP F2/3 with uncultured CB populations

The Gene Expression Omnibus dataset GSE125345 contains standardized RNA-Seq gene expression data of distinct hematopoietic cell states from uncultured cord blood. MEP, EryP, GMP, Gr, Mono and MEP populations were selected to be compared with the genes differentially expressed in TFEB^WT^ vs CTRL MEP F2/3 cells. The top 250 genes enriched in MEP F2/3 TFEB^WT^ and the top 250 genes enriched in MEP F2/3 CTRL were selected to be used as the reference signatures. Firstly, bar graphs were created by selecting the MEP F2/3 TFEB^WT^ reference signature, and calculating the number of scaled data that were above (>0) or below (<0) the mean for each population, corrected by the number of samples per population and 1,000 random permutations. Secondly, enrichment of both the TFEB^WT^ and CTRL MEP F2/3 reference signatures were estimated in each sample of the uncultured cord blood population using ssgsea() from the GSVA_1.30.0 R package.

### ATAC-seq processing and analysis

Transduced BFP^+^ LT-HSC: Library preparation for ATAC-Seq was performed on 1000-5000 cells with Nextera DNA Sample Preparation kit (Illumina), according to previously reported protocol(*41*). 4 ATAC-seq libraries were sequenced per lane in HiSeq 2500 System (Illumina) to generate paired-end 50-bp reads. Reads were mapped to hg38 using BWA (0.7.15) using default parameters. Duplicate reads, reads mapped to mitochondria, an ENCODE blacklisted region or an unspecified contig were removed (Encode Project Consortium, 2012). MACS (2.2.5) was used to call peaks in mapped reads. A catalogue of all peaks was obtained by concatenating all peaks and merging any overlapping peaks. Peaks were considered unique to one condition or another if they were present in at least 2 out of three replicates but not in the contrasting condition. Homer (4.11.1) was used to calculate enrichment of subsets of peaks using default parameters plus the catalogue of called peaks as background.

