## Supplemental Figures for "Dichotomous regulation of lysosomes by MYC and TFEB controls hematopoietic stem cell fate"

**Supplemental figure legends**

**Fig. S1.**

**A.** Sorting scheme for the indicated populations from CD34-column enriched human cord blood (hCB). **B.** Gene expression (RNAseq) of the indicated genes in uncultured (qLT-HSC) and 4d cultured (aLT-HSC) LT-HSC. n=3CB. **C.** Cell area was quantified by ImageJ/Fiji in images from Fig. 1C. n=3CB. Mann-Whitney test. **D.** Representative 3D projections of z-stack images from Fig. 1C. **E.** RNA-seq gene expression values of MITF, TFEB, TFE3 and TFEC in different hematopoietic cell populations isolated from CB (GSE12534). CQN: conditional quantile normalization. **F.** RNA-seq gene expression values of MITF, TFEB, TFE3 and TFEC in different hematopoietic populations isolated from bone marrow and/or peripheral blood (Hemaexplorer, http://servers.binf.ku.dk/bloodspot/, GSE17054, GSE19599, GSE11864, E-MEXP-1242). *P<0.05, **P<0.01, ***P<0.001. Unpaired t-test unless otherwise indicated.

**Fig. S2.**

**A.** RNA-seq gene expression values of MITF, TFEB, TFE3 and TFEC in HSC and progenitors (PROG) hematopoietic cell populations isolated from CB, BM and FL (GSE109093). **B**. RNA-seq gene expression values of MITF, TFEB, TFE3 and TFEC in different hematopoietic cell populations isolated from CB (GSE76234). **C,D,E.** Confocal analysis of TFEB and TFE3 expression in qLT-HSC and aLT-HSC (cultured for 48h). Graphs show the correlation of total (left) or nuclear (right) IntDen of TFEB and TFE3, and their percentage of cytoplasmic signal (E) measured in ImageJ/Fiji. Representative pictures are shown. Scale 2um. n=3CB, 311 cells. Mann-Whitney test. *P<0.05, **P<0.01, ***P<0.001.

**Fig. S3.**

**A**. Representative pictures from Fig. 1E. Scale 5um. n=3CB, 594 cells. **B.** qLT-HSC and cultured LT-HSC for 24h or 48h were stained with LysoSensor Green and analyzed by flow cytometry. n=3CB. **C**. Confocal analysis of MYC immunostaining in $MYC^{WT}$ and CTRL LT-HSC 3d post-transduction. Scale 2um. n=3CB, 270 cells. Mann-Whitney test. **D.** $MYC^{WT}$ and

CTRL LT-HSC were stained with MitoTracker Green and analyzed by flow cytometry 3d post-transduction. n=3CB. Paired t-test. **E,F.** GSEA analysis of the indicated gene sets in LT-HSC from Fig. 1F. FDRq-value<=0.05. n=3CB. **G.** Sorted LT-HSC were cultured for 4d in the presence of DMSO (control) or MYC inhibitor and stained with MitoTracker Green, CellROX Orange for flow cytometry analysis. n=3CB**. H.** Cell cycle analysis of LT-HSC from (G) by flow cytometry. n=3CB. *P<0.05, **P<0.01, ***P<0.001. Unpaired t-test unless otherwise indicated.

**Fig. S4.**

**A.** Confocal analysis of TFEB immunostaining in $TFEB^{WT}$-OE and CTRL LT-HSC 3d post-transduction. n=3CB, 323 cells. Mann-Whitney test. **B.** Differentially expressed genes (FDRq-value<=0.001) for LT-HSC from Fig. 2A. **C.** GSEA plot of lysosomal genes in LT-HSC from Fig. 2A. FDRq-value<=0.05. **D.** LT-HSC from Fig. 2A were subjected to ATAC-seq analysis at 6d post-transduction. Homer was used to calculate transcription factor recognition site motifs enriched in the set of peaks which were exclusive to $TFEB^{WT}$-OE LT-HSC relative to CTRL, using the catalogue of all peaks identified in any LT-HSC condition as a background. n=3CB. **E.** Logograms of MITF and TFEB frequency matrices from the Jaspar database. **F.** GSEA was used to calculate enrichment of the set of genes with a TFEB-OE specific chromatin accessibility peak within 2.5kb upstream and 0.5kb downstream of a GENCODE GRCh38 (https://www.ncbi.nlm.nih.gov/pubmed/30357393) TSS within a ranked list of genes by expression in $TFEB^{WT}$ vs CTRL LT-HSC (left) or in shTFEB vs shCTRL LT-HSC (right). FDRq-value<=0.05. n=3CB. **G.** Confocal analysis of Cathepsin D (CTSD) immunostaining in $TFEB^{WT}$-OE and CTRL LT-HSC 3d post-transduction. n=3CB, 323 cells. Mann-Whitney test. **H.** Representative 3D projections of z-stack images from Fig. 2F. **I.** RNA-seq expression of *TFRC* in LT-HSC from Fig. 1F (MYC-OE), Fig. 2A (TFEB-OE) and Fig. S5H (shTFEB). n=3 CB. **J.** Percentage of TfR1 signal overlapping with LAMP1 signal in pictures from Fig. 2F and analyzed in ImageJ/Fiji. n=3CB, 308 cells. Mann-Whitney test. **K.** Experimental scheme: sorted LT-HSC were cultured for 1d prior to transduction with lentiviral vectors expressing mCherry and a shRNA against Renilla (shCTRL) or LAMP1 (shLAMP1). $mCherry^{+}$ cells were sorted at 6d and analyzed for LAMP1 staining by confocal microscopy. n=3CB, 250 cells. Mann-Whitney test. **L.** TfR1 MFI analyzed by flow cytometry in LT-HSC from (K) at 3 and 6d post-transduction. n=3CB. Paired t-est. **M.** TfR1 MFI analyzed by flow cytometry in qLT-HSC and cultured LT-HSC for 24 and 48h. n=3CB. **N.** RNA-seq expression of *TFRC*, *MYC* and *TFEB* in CTRL LT-HSC at indicated time points. n=3CB. *P<0.05, **P<0.01, ***P<0.001. Unpaired t-test unless otherwise indicated.

**Fig. S5**

**A.** Gene expression (RNAseq) of indicated gene sets in LT-HSC from Fig. 2A. **B.** GSEA plots of indicated gene sets in LT-HSC from Fig. 2A and Fig. S5H. FDRq-value<=0.05. **C.** Enrichment map of gene sets with positive NES (red node) in $MYC^{WT}$ vs CTRL LT-HSC (280 gene sets FDRq-vale<=0.05) and shTFEB vs CTRL LT-HSC (130 gene sets FDRq-vale<=0.05) and negative NES (blue node) in $TFEB^{WT}$ vs CTRL LT-HSC (122 gene sets FDRq-vale<=0.05). Node size is proportional to the number of genes included in each gene set (minimum 10 genes/gene set). Grey edges indicate gene overlap. Clusters were automatically annotated using Autoannotate app in Cytoscape (grey circles). **D.** LT-HSC from Fig. 2A were stained with MitoTracker and CellROX and analyzed by flow cytometry at 3d post-transduction. n=3CB. **E.** Percentage of $BFP^{+}$ LT-HSC from Fig. 2A analyzed by flow cytometry at 3 to 6d post-transduction. n=3CB. **F.** LT-HSC from Fig. 2A were cultured for 1h with EdU and fixated for staining and flow cytometry analysis at 6d post-transduction. n=3CB. **G.** qPCR analysis of *TFEB* mRNA levels in Nalm6 cells transduced with 3 different shTFEB and one shCTRL lentiviral vectors. n=3 replicates. **H.** Experimental scheme: sorted LT-HSC were cultured for 1 day prior to transduction with lentiviral vectors expressing mCherry and a shRNA against Renilla (shCTRL) or TFEB (shTFEB). $mCherry^{+}$ cells were sorted at 6d for cell cycle analysis. n=3CB. **I.** LT-HSC from (H) were stained with Magic Red and analyzed by flow cytometry. n=3CB. *P<0.05, **P<0.01, ***P<0.001. Unpaired t-test unless otherwise indicated.

**Fig. S6.**

**A.** Samples tests are quiescent LT-HSC and the rows are tested gene-sets which are the top 100, 250 and 500 up or down regulated genes in $TFEB^{WT}$-OE vs CTRL LT-HSC or up and down regulated genes in $MYC^{WT}$-OE vs CTRL LT-HSC. Left: heatmap of GSVA scores. Right: Boxplot and strip chart showing higher enrichment score for the $TFEB^{WT}$-OE upregulated and $MYC^{WT}$-OE downregulated genes in quiescent LT-HSC. **B.** Graphs showing the enrichment score of MYC-OE and TFEB-OE upregulated genes (FDR q-value <=0.05) in 3 single-cell RNA-seq datasets(*26*,*27*,*28*). For more information on how the cell-cycle primed and non-primed cell clusters were identified, please see material and methods(*7*). **C.** Fold change in TfR1 membrane levels analyzed by flow cytometry in LT-HSC cultured for 4d with DMSO or MYC inhibitor. n=3CB. **D.** Experimental scheme: sorted LT-HSC were cultured for 1 day prior to transduction with lentiviral vectors expressing mCherry and a shRNA against Renilla (shCTRL) or TFRC (shTFRC). mCherry$^+$ cells were sorted at 3 and 6d for TfR1 MFI analysis by flow cytometry. n=3CB. **E.** LT-HSC from (D) were stained with Ki67 and Hoechst and analyzed by flow cytometry. n=3CB. **F,G,H.** LT-HSC from Fig. 3A were stained with LysoSensor Green (F), Magic Red (G) and TfR1 antibody (H) and analyzed by flow cytometry. n=3CB. Paired t-test. **I**. Number of cells per colony was measured by flow cytometry from CFC assays from Fig. 3A. n=3CB. **J.** Cells pooled from CFC colonies from Fig. 3A were stained for the indicated lineage markers and analyzed by flow cytometry. n=3CB. **K,L,M,N**. Individual LT-HSC from Fig. 3A were sorted and plated on stromal cell layers pre-seeded in 96-well plates (SCS assay) and after 16-17 days the number of colonies was scored by microscopy (K). Each well containing a colony was stained with lineage markers and analyzed by flow cytometry(L,M,N). n=3CB *P<0.05, **P<0.01, ***P<0.001. Unpaired t-test unless otherwise indicated.

**Fig. S7.**

**A,B.** RNA-seq normalized counts for *TFEB* and *MYC* mRNA in the indicated cell populations sorted from hCB using the membrane markers listed in the table (B) and in Fig. S1A. n=3CB. **C.** Experimental scheme: LT-HSC were cultured for 1 day prior transduction with lentiviral vectors expressing BFP and one of the following genes: CTRL or TFEB$^{WT}$. BFP$^+$ cells were sorted at day 3 and plated for CFC and SCS assays. **D,E,F.** SCS assay: each well containing a colony from (C) was stained with lineage markers and analyzed by flow cytometry. Paired t-test. n=3CB. **G.** Experimental scheme: sorted ST-HSC following the sorting scheme described in Fig. S1A were cultured for 1d prior transduction with lentiviral vectors expressing BFP and one of the following genes: CTRL or TFEB$^{WT}$. BFP$^+$ cells were sorted at day 3 and plated for CFC and SCS assays. **H,I**. CFC assays with ST-HSC from (G). Colonies were scored under the microscope (H) and cells pooled from colonies were stained for the indicated lineage markers and analyzed by flow cytometry (I). n=3CB. **J,K.** SCS assay with ST-HSC from (G). The number of colonies was scored under the microscope (J) and each well containing a colony was stained with lineage markers and analyzed by flow cytometry (K). n=3CB. Paired t-test. **L.** Experimental scheme: sorted LT-HSC were cultured for 1d prior to transduction with lentiviral vectors expressing mCherry and a shRNA against Renilla (shCTRL) or TFEB (shTFEB). mCherry$^+$ cells were sorted at 3d for SCS assays. **M,N,O.** SCS assay with LT-HSC from (L). The number of colonies was scored under the microscope (M) and each well containing a colony was stained with lineage markers and analyzed by flow cytometry (N,O). n=3CB. **P.** Experimental scheme: sorted LT-HSC were cultured for 1d prior to transduction with lentiviral vectors expressing mCherry and a shRNA against Renilla (shCTRL) or LAMP1 (shLAMP1). mCherry$^+$ cells were sorted at 6d for CFC assays. **Q.** CFC assay with LT-HSC from (P). Colonies were scored under the microscope and cells pooled from colonies were stained for the indicated lineage markers and analyzed by flow cytometry. n=3CB. **R.** Experimental scheme: sorted LT-HSC were cultured for 1d prior to transduction with lentiviral vectors expressing mCherry and a shRNA against Renilla (shCTRL) or TFRC (shTFRC). mCherry$^+$ cells were sorted at 6d for CFC assays. **S.** CFC assay with LT-HSC from (R). Colonies were scored under the microscope and cells pooled from colonies were stained for the indicated lineage markers and analyzed by flow cytometry. n=3CB. **T.** Experimental scheme: MEP F1 and MEP F2/3 were sorted from CD34-enriched hCB following sorting scheme displayed in Fig. 1A and cultured for 1d prior to transduction with lentivirus expressing BFP and one of the following genes: CTRL or TFEB$^{WT}$. BFP$^+$ cells were sorted at day 3d and plated for CFC and SCS assays. **V.** TfR1 expression in MEP F2/3 cells from (T) at 3 and 6d post-transduction. n=3-6CB. *P<0.05, **P<0.01, ***P<0.001. Unpaired t-test unless otherwise indicated.

**Fig. S8.**

**A. B, C.** CFC assays with MEP F1 and F2/3 cells from Fig. S7T. Colonies were scored 10-11 days after plating and cells were pooled for flow cytometry analysis. n=3CB. **D,E,F,G,H,I**. SCS assays with MEP F1 and F2/3 cells from Fig. S7T. Individual wells containing a colony were scored by microscopy, stained for lineage markers and analyzed by flow cytometry. n=3CB. **J.** RNA-seq expression of *TFEB* and *MYC* in MEP F2/3 cells from Fig. S7T. n=2-3CB. **K,L**. GSEA plots and expression of indicated gene sets in MEP F2/3 cells from Fig. S7T. n=2-3CB. **M,N**. GSVA analysis with the 250 most upregulated genes between TFEB$^{WT}$-OE and CTRL

MEP F2/3 cells in the indicated populations sorted from hCB. *P<0.05, **P<0.01, ***P<0.001. Unpaired t-test unless otherwise indicated.

**Fig. S9.**

**A.** TFEB protein domains and sequence conservation of the bHLH domain (adapted from Puertollano et al., EMBO 2018)(Puertollano et al., 2018). Multiple sequence alignment highlights the conservation of critical domains between human TFEB, TFE3, MITF and TFEC proteins. S142A (in blue) is the mutation introduced to create $TFEB^{S142A}$. H240R and I243N (in red) are the two mutations introduced to create $TFEB^{H240R-I243N}$. **B.** Experimental scheme: $CD34^{+}CD38^{-}$ cells were sorted from CD34-enriched hCB and transduced *in vitro* using lentiviral vectors expressing BFP and one of the following genes: CTRL, $TFEB^{WT}$, $TFEB^{S142A}$ or $TFEB^{H240R-I243N}$. At day 8 post-transduction $BFP^{+}$ cells were sorted for the indicated analysis. **C.** $CD34^{+}CD38^{-}$ cells from (B) were fixated on slides and stained with TFEB antibody and DAPI solution. Representative images captured by a Zeiss LSM700 Confocal (oil, 63x/1.4NA, inverted, Zen 2012 software) are shown. Scale 5um. **D.** qPCR of *TFEB* expression in Nalm6 cells transduced with CTRL, $TFEB^{WT}$, $TFEB^{S142A}$ or $TFEB^{H240R-I243N}$ lentivirus 5d post-transduction. n=3CB. **E.** $CD34^{+}CD38^{-}$ cells from (B) were stained with CytoID and analyzed by flow cytometry at 8d post-transduction. n=3CB. **F.** $CD34^{+}CD38^{-}$ cells from (B) were analyzed by flow cytometry at the indicated time points and the fold change of %$BFP^{+}$ cells was calculated. n=2CB. **G**. GSVA analysis showing enrichment of KEGG lysosome pathway in HSC (LT-HSC) vs different subpopulations of MPP (ST-HSC) isolated from CB (GSE76234)(*21*). **H.** GSEA analysis showing enrichment of the KEGG lysosome pathway in HSC1 (LT-HSC) vs MPP (ST-HSC) isolated from CB (GSE42414)(*30*) (FDR q value <= 0.05). **I.** GSEA plots of indicated gene sets in 4d cultured LT- vs ST-HSC using RNA-seq data. FDRq-value<=0.05. n=2-3CB. **J.** Expression of indicated genes in $TFEB^{WT}$ vs CTRL ST-HSC from Fig. 4I. n=2CB**.** **K.** NES values for gene sets differentially enriched in $TFEB^{WT}$ vs CTRL ST-HSC from Fig. 4I. n=2CB**. L.** ST-HSC from Fig. 4I were cultured for 1h with EdU and fixated for staining and flow cytometry analysis. n=3CB. **M.** ST-HSC were cultured for 4d with DMSO (control) or MYC inhibitor, stained with MitoTracker Green, CellROX Orange and TfR1 and analyzed by flow cytometry. n=3CB**.** *P<0.05, **P<0.01, ***P<0.001. Unpaired t-test unless otherwise indicated.

**Fig. S10.**

**A.** Experimental scheme: LT-HSC were cultured for 1d prior to transduction with lentiviral vectors expressing BFP and CTRL or $TFEB^{WT}$. $BFP^+$ cells were sorted at day 3d for serial CFC assays. **B,C,D**. Colonies were scored under the microscope and number of cells analyzed by flow cytometry. $2^o$ and $3^o$ CFC assays were performed following the same protocol and timing. n=3-6CB. Paired t-test. **E,F.** Sorted LT-HSC were cultured for 1d prior to transduction with lentiviral vectors expressing mCherry and a shRNA against Renilla (shCTRL) or TFEB (shTFEB). $mCherry^+$ cells were sorted at 6d for serial CFC assays as in (A). Colonies were scored under the microscope and number of cells analyzed by flow cytometry. n=3CB. Paired t-test. **G.** From Fig. 5A: Percentage of total $CD45^+$ human cells at 17 weeks of engraftment in male NSG-W41 mice. **H.** From Fig. 5A: Percentage of total $CD45^+$ human cells at 4 weeks of engraftment in male NSG mice. **I.** From Fig. 5A: LOG2 ratio of $\%BFP^+$ in $CD45^+$ human cells at 4 weeks of engraftment in male NSG mice vs input (3d post-transduction). **J.** From Fig. 5A: Percentage of total $CD45^+$ human cells at 17 weeks of engraftment in female NSG mice. **K.** From Fig. 5A: LOG2 ratio of $\%BFP^+$ in $CD45^+$ human cells at 17 weeks of engraftment in female NSG mice vs input (3d post-transduction). **L.M.** From Fig. 5A: Table showing the $CD45^+BFP^+$cell doses injected in secondary NSG-GM3 mice, the number of injected mice per cell dose and the number of engrafted mice per cell dose and condition, at 8 weeks post-xenotransplantation. This data and the ELDA software were used to calculate the stem cell frequency shown in Fig. 5C and summarized in (M). Each *in vivo* experiment was performed with 3 independent CB pools and each dot represents an injected mouse (4-5 mice/CB). (RF: right femur, injected bone; BM: left femur and right and left tibiae; SP: spleen). *P<0.05, **P<0.01, ***P<0.001. Unpaired t-test unless otherwise indicated.

**Fig. S11.**

**A.** From Fig. 5D: Percentage of total $CD45^+$ human cells at 17w of engraftment in male NSG-W41 mice. **B.** From Fig. 5D: Percentage of total $CD45^+$ human cells at 4w of engraftment in male NSG mice. **C.** From Fig. 5D: Fold change of $\%mCherry^+$ in $CD45^+$ human cells at 4w of engraftment in male NSG mice vs input (3d post-transduction). **D.** From Fig. 5D: Percentage of total $CD45^+$ human cells at 17w of engraftment in female NSG mice. **E.** From Fig. 5D: Fold change of %mCherry$^+$ in CD45$^+$ human cells at 17w of engraftment in female NSG mice vs input (3d post-transduction). **F,G.** From Fig. 5D: Table showing the CD45$^+$mCherry$^+$cell doses injected in secondary NSG-GM3 mice, the number of injected mice per cell dose and the number of engrafted mice per cell dose and condition, at 8w post-xenotransplantation. This data and the ELDA software were used to calculate the stem cell frequency shown in Fig. 5F and summarized in (G). **H.** From Fig. 5A: Percentage of CD33$^+$ from total CD45$^+$BFP$^+$ cells analyzed by flow cytometry at 4w of xenotransplantation in NSG mice. **I.** From Fig. 5A: Percentage of CD33$^+$ from total CD45$^+$BFP$^+$ cells analyzed by flow cytometry at 17w of xenotransplantation in NSG-W41 mice. **J.** From Fig. 5D: Percentage of GlyA$^+$ from total CD45$^+$mCherry$^+$ cells analyzed by flow cytometry at 17w of xenotransplantation in NSG-W41 mice. Each *in vivo* experiment was performed with 3 independent CB pools and each dot represents an injected mouse (4-5 mice/CB). (RF: right femur, injected bone; BM: left femur and right and left tibiae; SP: spleen). *P<0.05, **P<0.01, ***P<0.001. Unpaired t-test unless otherwise indicated.

**Fig. S12.**

**A.** Experimental scheme: LT-HSC isolated from hCB were cultured for 4d and stained with LysoTracker Blue to sort LysoH and LysoL LT-HSC that were intrafemorally injected into 8-10w old male NSG-W41 mice. NSG-W41 mice were sacrificed at 4 and 17w post-xenotransplantation for analysis. CD45$^+$ human cells were sorted from primary NSG-W41 mice at 17w for secondary limiting dilution assays (LDA) in NSG-GM3 mice. NSG-GM3 mice were sacrificed at 8w post-xenotransplantation for stem cell frequency analysis. **B.** From (A): Percentage of total CD45$^+$ human cells at 4 and 17w of engraftment in male NSG-W41 mice. **C.** From (A): Percentage of GlyA$^+$ from total hCD45$^+$ cells analyzed by flow cytometry at 17w of xenotransplantation in NSG-W41 mice. **D,E.** From (A): Table showing the hCD45$^+$ cell doses injected in secondary NSG-GM3 mice, the number of injected mice per cell dose and the number of engrafted mice per cell dose and condition, at 8w post-xenotransplantation. This data and the ELDA software were used to calculate the stem cell frequency shown in Fig. 5G and summarized in (E). Each *in vivo* experiment was performed with 3 independent CB pools and each dot represents an injected mouse (4-5 mice/CB). (RF: right femur, injected bone; BM: left femur and right and left tibiae). *P<0.05, **P<0.01, ***P<0.001. Unpaired t-test unless otherwise indicated. **F.** Graphical abstract: Lysosomes are key signaling hubs for demandadapted regulation of LT-HSC fate. Lysosomes are dichotomously orchestrated by TFEB and MYC, which are instrumental to balance catabolic and anabolic reactions required to fuel divergent LT-HSC fate choices. TFEB localizes in the nucleus of quiescent LT-HSC where it induces the expression of endolysosomal genes required for membrane receptor endocytosis and lysosomal degradation. TFEB overexpression in LT-HSC reduces the membrane levels of TfR1 and imposes a quiescent low metabolic state preventing LT-HSC from activation and exhaustion while promoting self-renewal. On the other hand, MYC counteracts TFEB function and drives LT-HSC activation by downregulating TFEB-regulated lysosomal program and TfR1 degradation. This tag and war between TFEB and MYC over lysosome control also regulates LT-HSC lineage commitment. Induction of TFEB-mediated lysosomal activity imposes myeloid program and suppresses erythroid differentiation in LT-HSC and downstream progenitors.

**Supplemental information**

**Table S1.** Quiescent vs Activated LT-HSC RNAseq

**Table S2.** $MYC^{WT}$ vs CTRL LT-HSC RNAseq

**Table S3.** $TFEB^{WT}$ vs CTRL LT-HSC RNAseq and pathway enrichment analysis

**Table S4.** $TFEB^{WT}$ vs CTRL LT-HSC ATAC-seq peaks

**Table S5.** shTFEB vs shCTRL LT-HSC RNAseq

**Table S6.** $TFEB^{WT}$ vs CTRL MEP F2/3 RNAseq

**Table S7**. Quiescent LT- vs ST-HSC RNA-seq

**Table S8.** Activated LT- vs ST-HSC RNA-seq

**Table S9.** $TFEB^{WT}$ vs CTRL ST-HSC RNAseq

**Movie S1.** Short and long time to division.

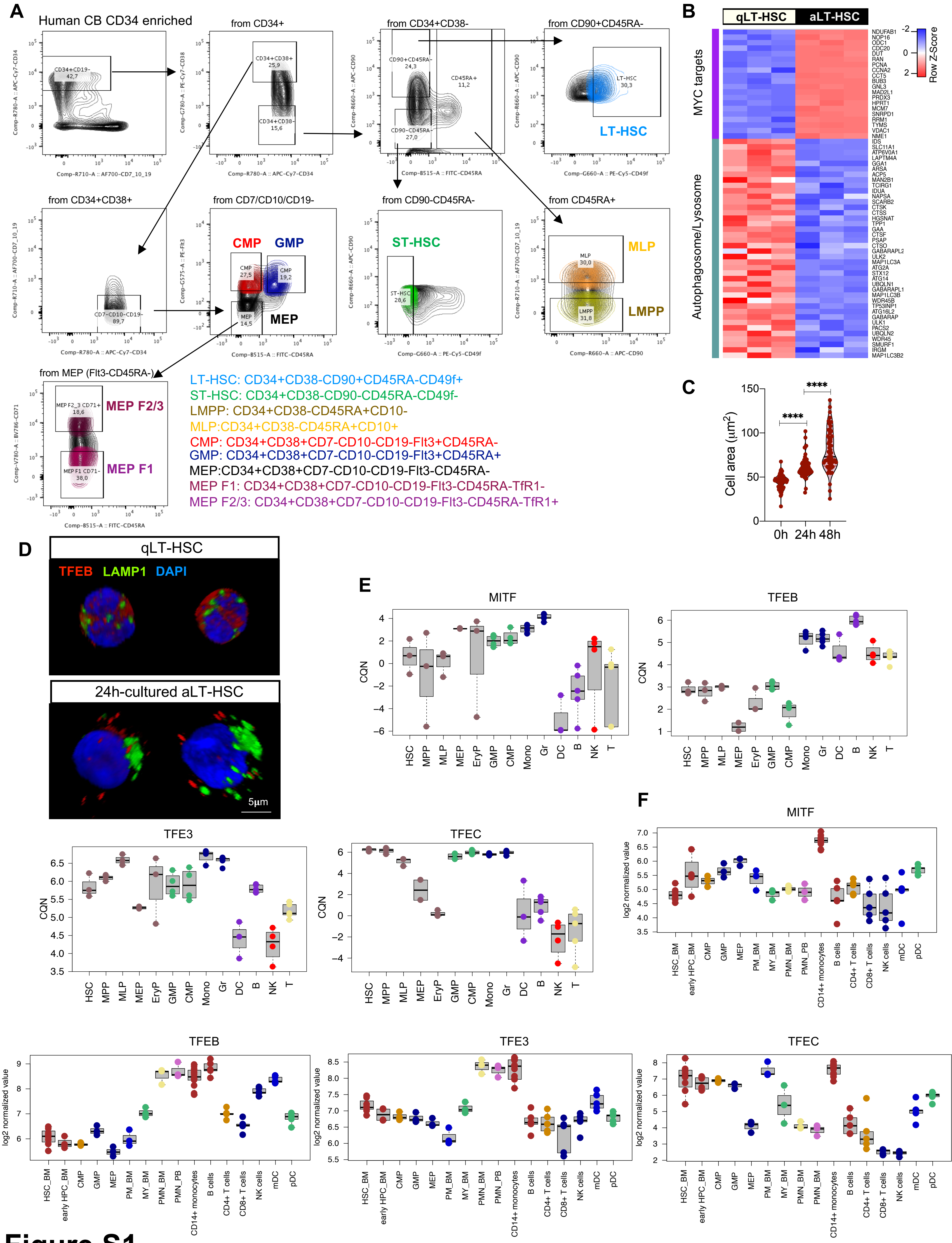

LT-HSC: CD34+CD38-CD90+CD45RA-CD49f+
ST-HSC: CD34+CD38-CD90-CD45RA-CD49f-
LMPP: CD34+CD38-CD45RA+CD10-
MLP:CD34+CD38-CD45RA+CD10+
CMP: CD34+CD38+CD7-CD10-CD19-Flt3+CD45RA-
GMP: CD34+CD38+CD7-CD10-CD19-Flt3+CD45RA+
MEP:CD34+CD38+CD7-CD10-CD19-Flt3-CD45RA-
MEP F1: CD34+CD38+CD7-CD10-CD19-Flt3-CD45RA-TfR1-
MEP F2/3: CD34+CD38+CD7-CD10-CD19-Flt3-CD45RA-TfR1+


### Figure S1

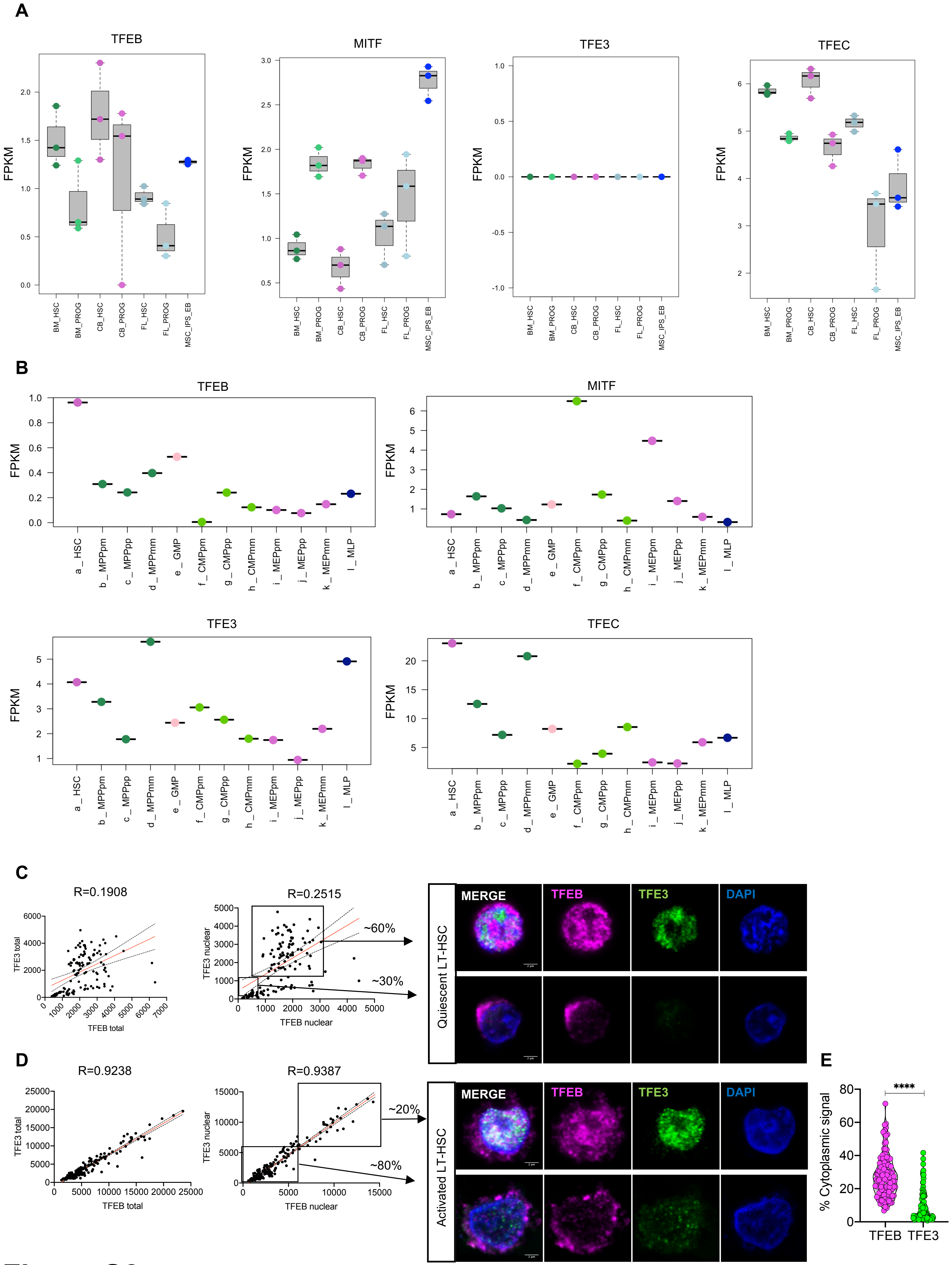
A
TFEB
MITF
TFE3
TFEC
FPKM
BM_HSC
BM_PROG
CB_HSC
CB_PROG
FL_HSC
FL_PROG
MSC_IPS_EB
B
TFEB
MITF
TFE3
TFEC
FPKM
a_HSC
b_MPPpm
c_MPPpp
d_MPPmm
e_GMP
f_CMPpm
g_CMPpp
h_CMPmm
i_MEPpm
j_MEPpp
k_MEPmm
l_MLP
C
R=0.1908
TFE3 total
TFEB total
R=0.2515
TFE3 nuclear
TFEB nuclear
~60%
~30%
Quiescent LT-HSC
MERGE
TFEB
TFE3
DAPI
D
R=0.9238
TFE3 total
TFEB total
R=0.9387
TFE3 nuclear
TFEB nuclear
~20%
~80%
Activated LT-HSC
MERGE
TFEB
TFE3
DAPI
E
% Cytoplasmic signal
****
TFEB
TFE3

Figure S2

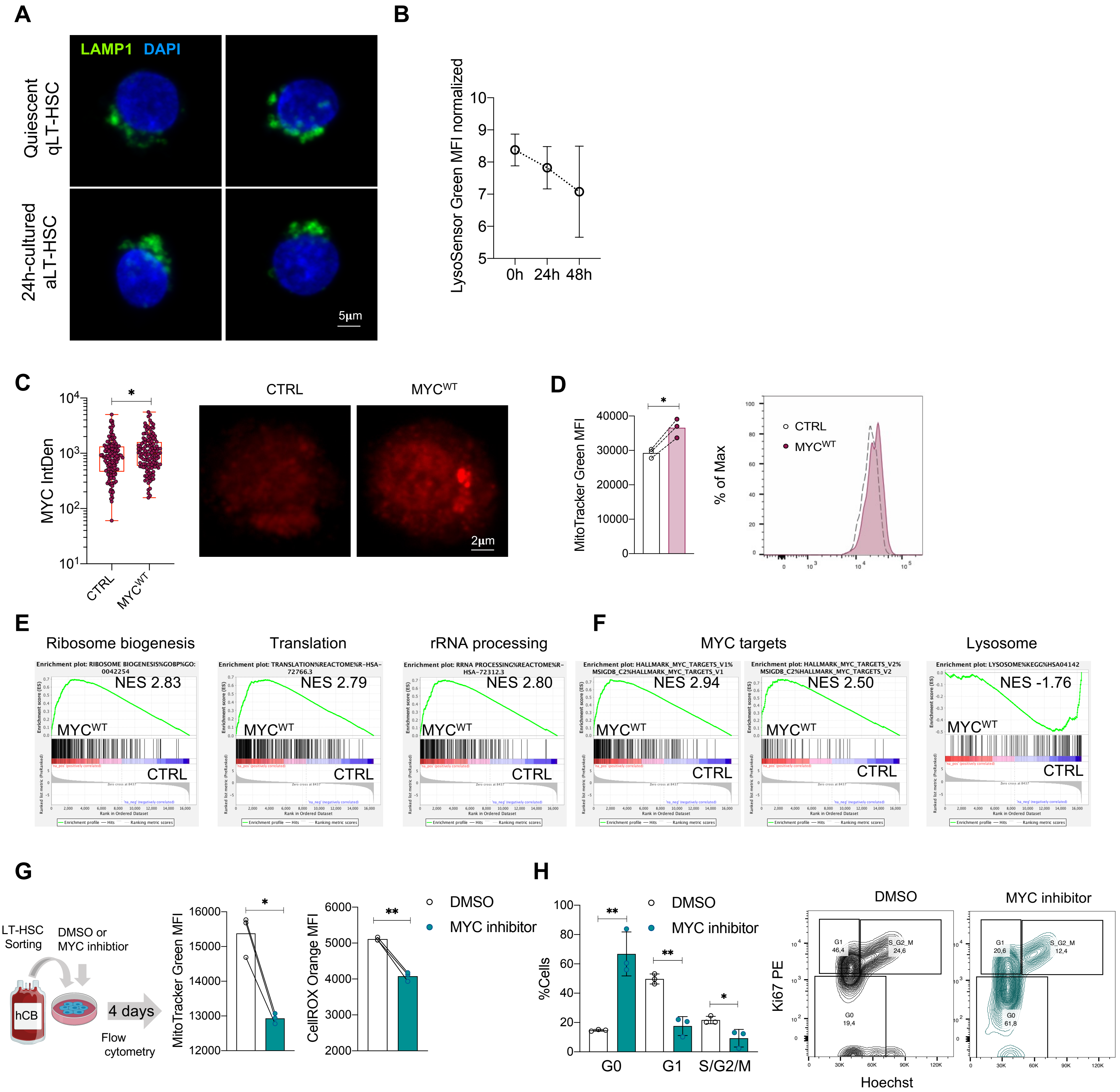
A
LAMP1 DAPI
Quiescent qLT-HSC
24h-cultured aLT-HSC
5µm
B
LysoSensor Green MFI normalized
0h 24h 48h
C
MYC IntDen
CTRL
MYCWT
2µm
D
MitoTracker Green MFI
% of Max
CTRL
MYCWT
E
Ribosome biogenesis
NES 2.83
Translation
NES 2.79
rRNA processing
NES 2.80
F
MYC targets
NES 2.94
NES 2.50
Lysosome
NES -1.76
G
LT-HSC Sorting
DMSO or MYC inhibtior
hCB
4 days
Flow cytometry
MitoTracker Green MFI
CellROX Orange MFI
DMSO
MYC inhibitor
H
%Cells
G0
G1
S/G2/M
DMSO
MYC inhibitor
Ki67 PE
Hoechst
G1 46,4
S_G2_M 24,6
G0 19,4
G1 20,6
S_G2_M 12,4
G0 61,8

### Figure S3

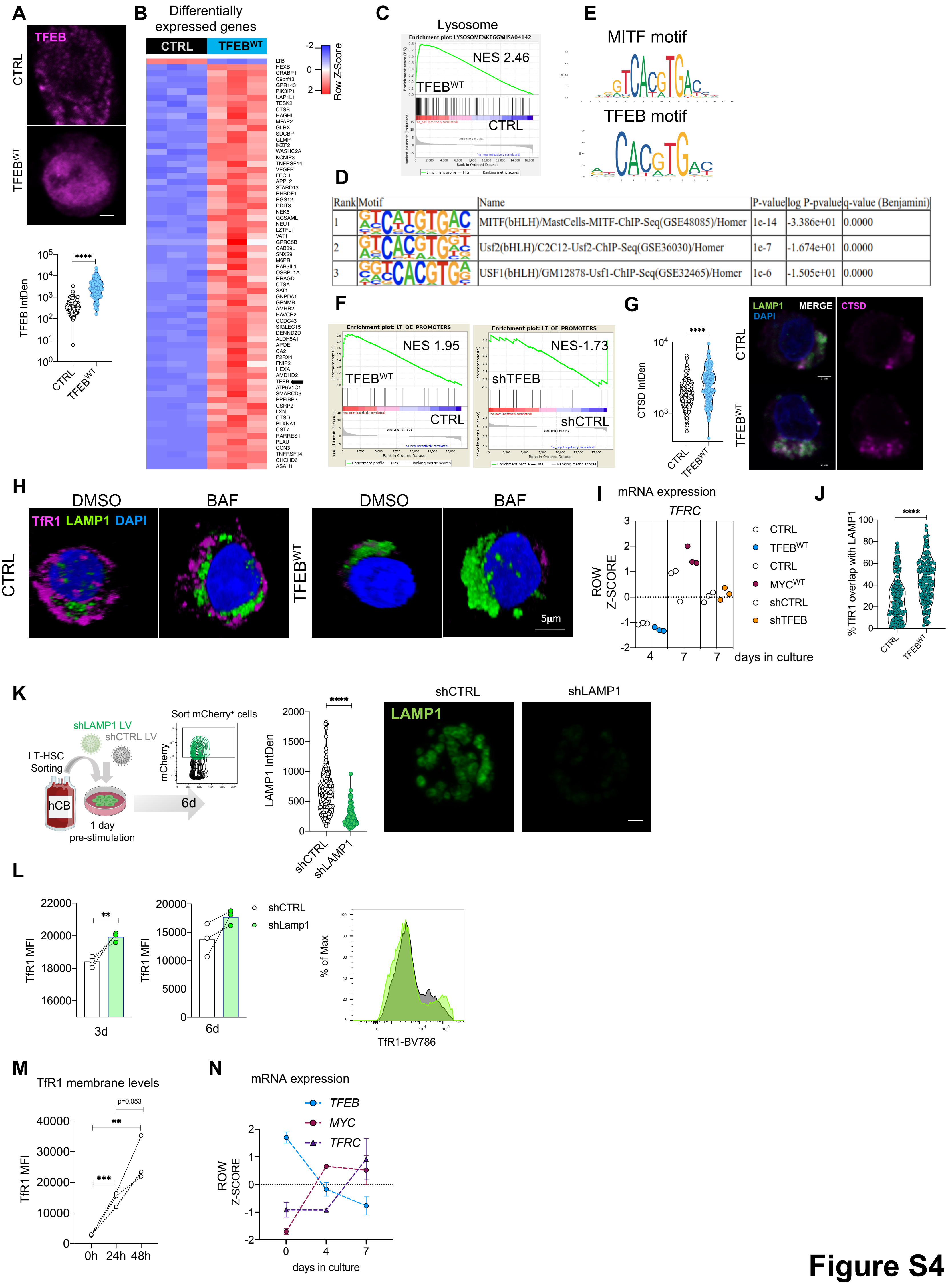
A
TFEB
CTRL
TFEB^WT
TFEB IntDen
****
B
Differentially expressed genes
CTRL
TFEB^WT
Row Z-Score
C
Lysosome
Enrichment plot: LYSOSOME%KEGG%HSA04142
NES 2.46
TFEB^WT
CTRL
D
Rank | Motif | Name | P-value | log P-pvalue | q-value (Benjamini)
1 | MITF(bHLH)/MastCells-MITF-ChIP-Seq(GSE48085)/Homer | 1e-14 | -3.386e+01 | 0.0000
2 | Usf2(bHLH)/C2C12-Usf2-ChIP-Seq(GSE36030)/Homer | 1e-7 | -1.674e+01 | 0.0000
3 | USF1(bHLH)/GM12878-Usf1-ChIP-Seq(GSE32465)/Homer | 1e-6 | -1.505e+01 | 0.0000
E
MITF motif
TFEB motif
F
Enrichment plot: LT_OE_PROMOTERS
NES 1.95
TFEB^WT
CTRL
NES-1.73
shTFEB
shCTRL
G
CTSD IntDen
****
LAMP1
DAPI
MERGE
CTSD
CTRL
TFEB^WT
H
DMSO
BAF
TfR1 LAMP1 DAPI
CTRL
TFEB^WT
5μm
I
mRNA expression
TFRC
ROW Z-SCORE
days in culture
CTRL
TFEB^WT
CTRL
MYC^WT
shCTRL
shTFEB
J
****
%TfR1 overlap with LAMP1
CTRL
TFEB^WT
K
shLAMP1 LV
shCTRL LV
LT-HSC Sorting
hCB
1 day pre-stimulation
Sort mCherry⁺ cells
mCherry
6d
LAMP1 IntDen
****
shCTRL
shLAMP1
LAMP1
L
TfR1 MFI
**
3d
6d
shCTRL
shLamp1
% of Max
TfR1-BV786
M
TfR1 membrane levels
TfR1 MFI
p=0.053
**
***
0h
24h
48h
N
mRNA expression
TFEB
MYC
TFRC
ROW Z-SCORE
days in culture

Figure S4

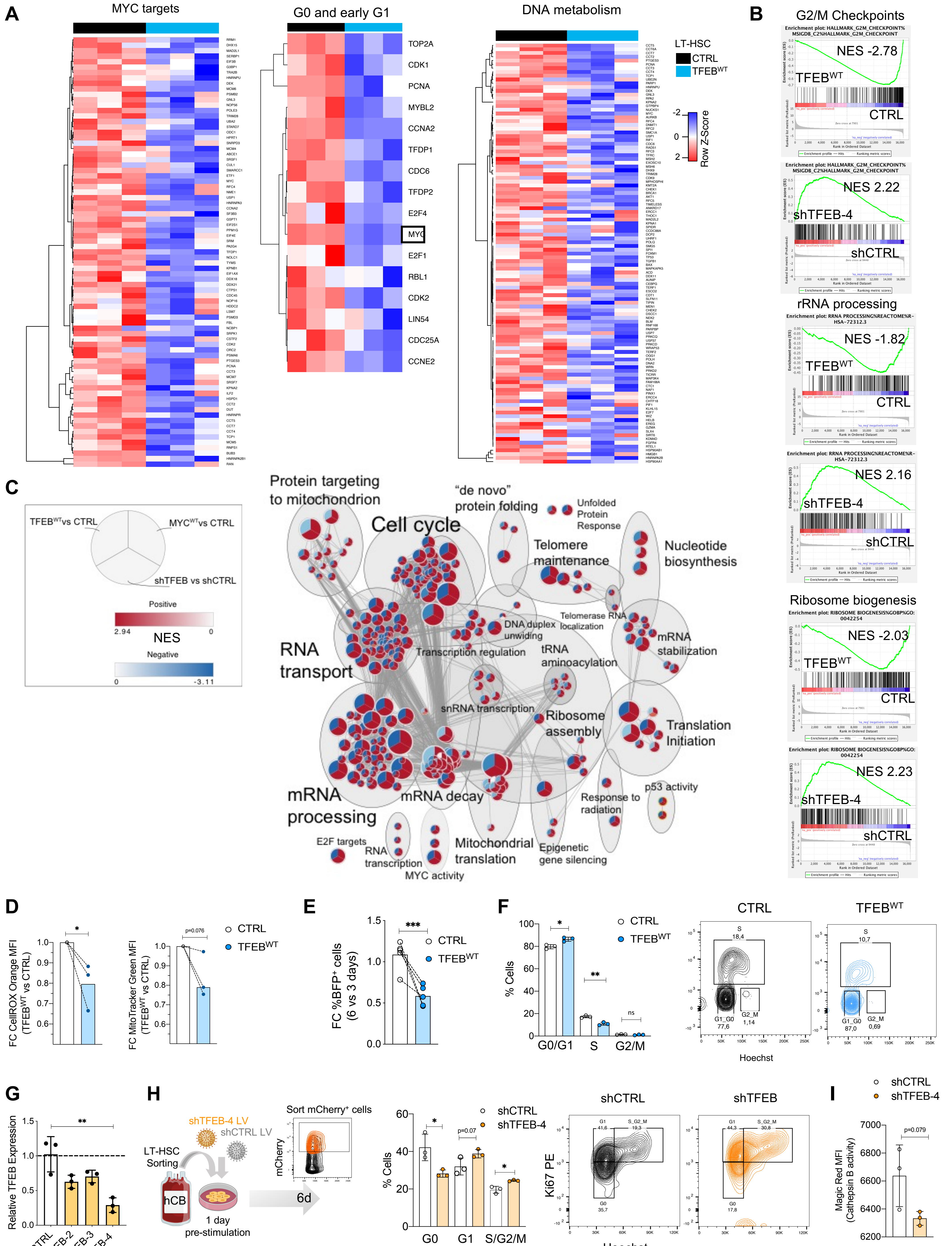
A
MYC targets
G0 and early G1
DNA metabolism
TOP2A
CDK1
PCNA
MYBL2
CCNA2
TFDP1
CDC6
TFDP2
E2F4
MYC
E2F1
RBL1
CDK2
LIN54
CDC25A
CCNE2
LT-HSC
CTRL
TFEB^WT
Row Z-Score
-2
0
2
B
G2/M Checkpoints
NES -2.78
TFEB^WT
CTRL
NES 2.22
shTFEB-4
shCTRL
rRNA processing
NES -1.82
TFEB^WT
CTRL
NES 2.16
shTFEB-4
shCTRL
Ribosome biogenesis
NES -2.03
TFEB^WT
CTRL
NES 2.23
shTFEB-4
shCTRL
C
TFEB^WT vs CTRL
MYC^WT vs CTRL
shTFEB vs shCTRL
Positive
2.94
NES
0
Negative
0
-3.11
Protein targeting to mitochondrion
Cell cycle
"de novo" protein folding
Unfolded Protein Response
Telomere maintenance
Nucleotide biosynthesis
DNA duplex unwinding
Telomerase RNA localization
mRNA stabilization
RNA transport
Transcription regulation
tRNA aminoacylation
snRNA transcription
Ribosome assembly
Translation Initiation
mRNA processing
mRNA decay
Response to radiation
p53 activity
E2F targets
RNA transcription
MYC activity
Mitochondrial translation
Epigenetic gene silencing
D
FC CellROX Orange MFI (TFEB^WT vs CTRL)
FC MitoTracker Green MFI (TFEB^WT vs CTRL)
p=0.076
CTRL
TFEB^WT
E
FC %BFP+ cells (6 vs 3 days)
***
CTRL
TFEB^WT
F
% Cells
CTRL
TFEB^WT
G0/G1
S
G2/M
ns
CTRL
TFEB^WT
S 18,4
G1_G0 77,6
G2_M 1,14
S 10,7
G1_G0 87,0
G2_M 0,69
Hoechst
G
Relative TFEB Expression
shCTRL
shTFEB-2
shTFEB-3
shTFEB-4
H
shTFEB-4 LV
shCTRL LV
LT-HSC Sorting
hCB
1 day pre-stimulation
Sort mCherry+ cells
mCherry
6d
% Cells
shCTRL
shTFEB-4
p=0.07
G0
G1
S/G2/M
shCTRL
shTFEB
Ki67 PE
G1 41,6
S_G2_M 19,3
G0 35,7
G1 44,3
S_G2_M 30,8
G0 17,8
Hoechst
I
shCTRL
shTFEB-4
Magic Red MFI (Cathepsin B activity)
p=0.079

Figure S5

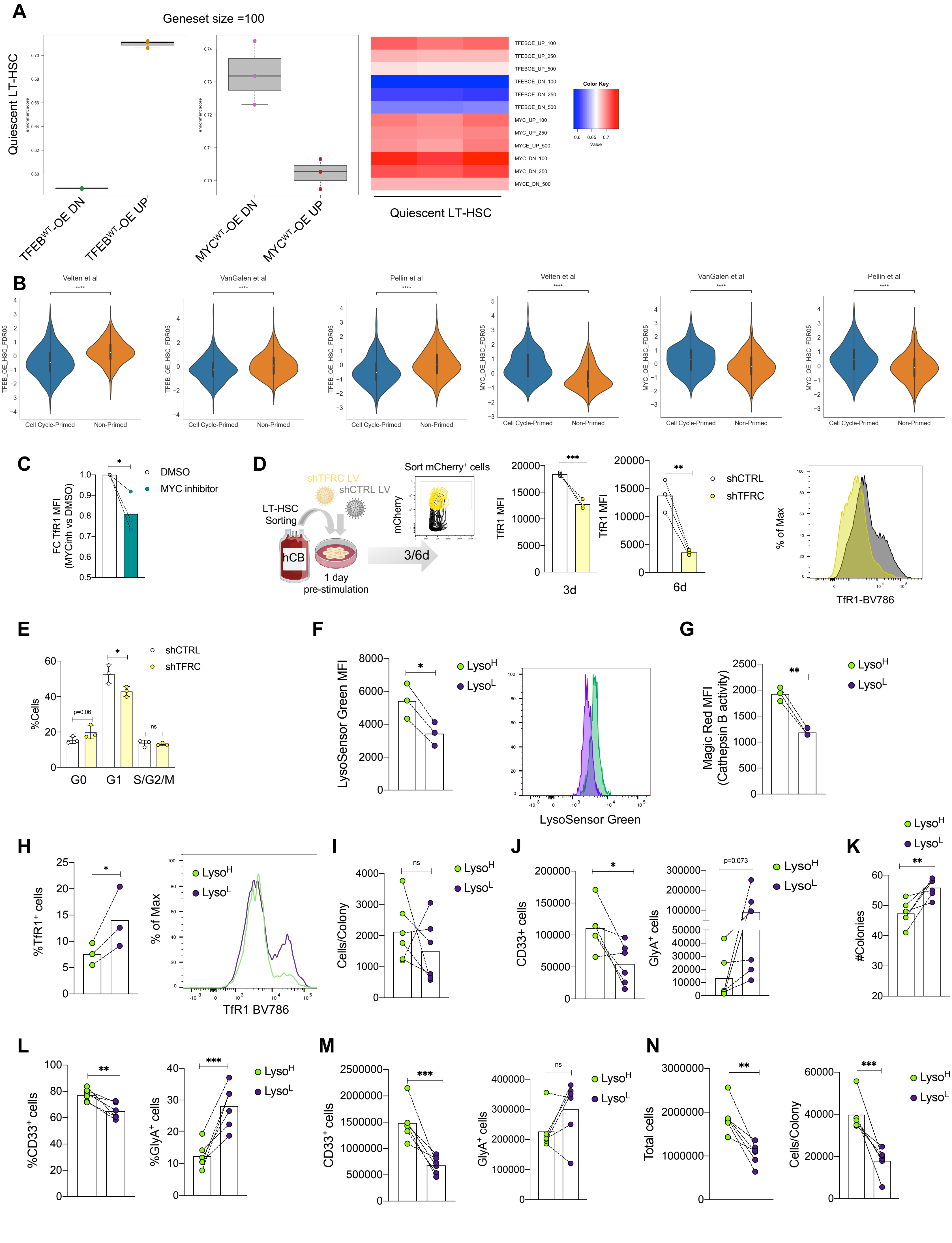
A
Geneset size =100
Quiescent LT-HSC
TFEB^WT-OE DN
TFEB^WT-OE UP
MYC^WT-OE DN
MYC^WT-OE UP
TFEBOE_UP_100
TFEBOE_UP_250
TFEBOE_UP_500
TFEBOE_DN_100
TFEBOE_DN_250
TFEBOE_DN_500
MYC_UP_100
MYC_UP_250
MYCE_UP_500
MYC_DN_100
MYC_DN_250
MYCE_DN_500
Color Key
Value
B
Velten et al
VanGalen et al
Pellin et al
TFEB_OE_HSC_FDR05
MYC_OE_HSC_FDR05
Cell Cycle-Primed
Non-Primed
C
FC TfR1 MFI (MYCinh vs DMSO)
DMSO
MYC inhibitor
D
shTFRC LV
shCTRL LV
LT-HSC Sorting
hCB
1 day pre-stimulation
3/6d
Sort mCherry+ cells
mCherry
TfR1 MFI
3d
6d
shCTRL
shTFRC
% of Max
TfR1-BV786
E
%Cells
G0
G1
S/G2/M
p=0.06
ns
F
LysoSensor Green MFI
Lyso^H
Lyso^L
LysoSensor Green
G
Magic Red MFI (Cathepsin B activity)
H
%TfR1+ cells
% of Max
TfR1 BV786
I
Cells/Colony
J
CD33+ cells
GlyA+ cells
p=0.073
K
#Colonies
L
%CD33+ cells
%GlyA+ cells
M
CD33+ cells
GlyA+ cells
N
Total cells
Cells/Colony


**Figure S6**

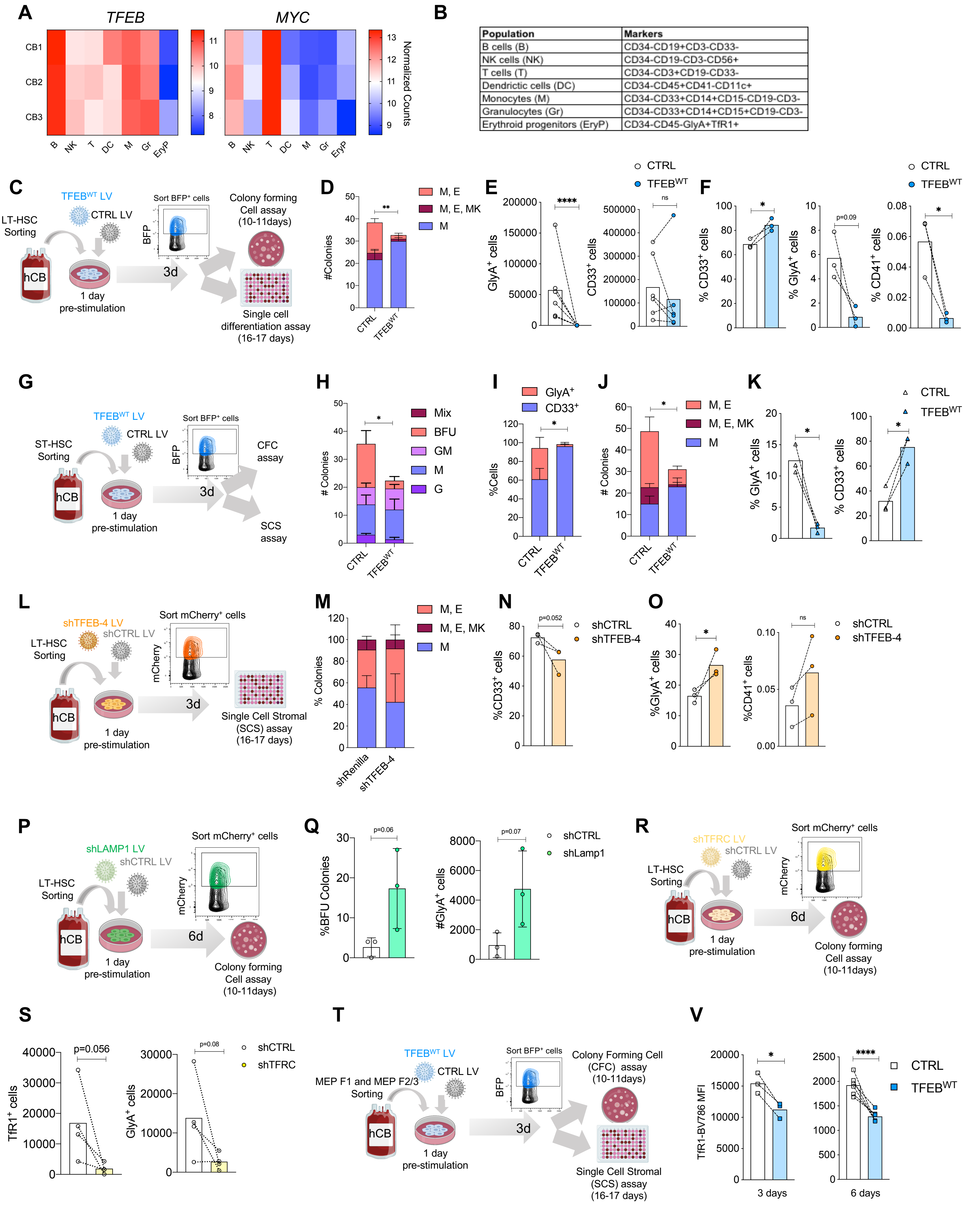


| Population | Markers |
|---|---|
| B cells (B) | CD34-CD19+CD3-CD33- |
| NK cells (NK) | CD34-CD19-CD3-CD56+ |
| T cells (T) | CD34-CD3+CD19-CD33- |
| Dendrictic cells (DC) | CD34-CD45+CD41-CD11c+ |
| Monocytes (M) | CD34-CD33+CD14+CD15-CD19-CD3- |
| Granulocytes (Gr) | CD34-CD33+CD14+CD15+CD19-CD3- |
| Erythroid progenitors (EryP) | CD34-CD45-GlyA+TfR1+ |

### Figure S7

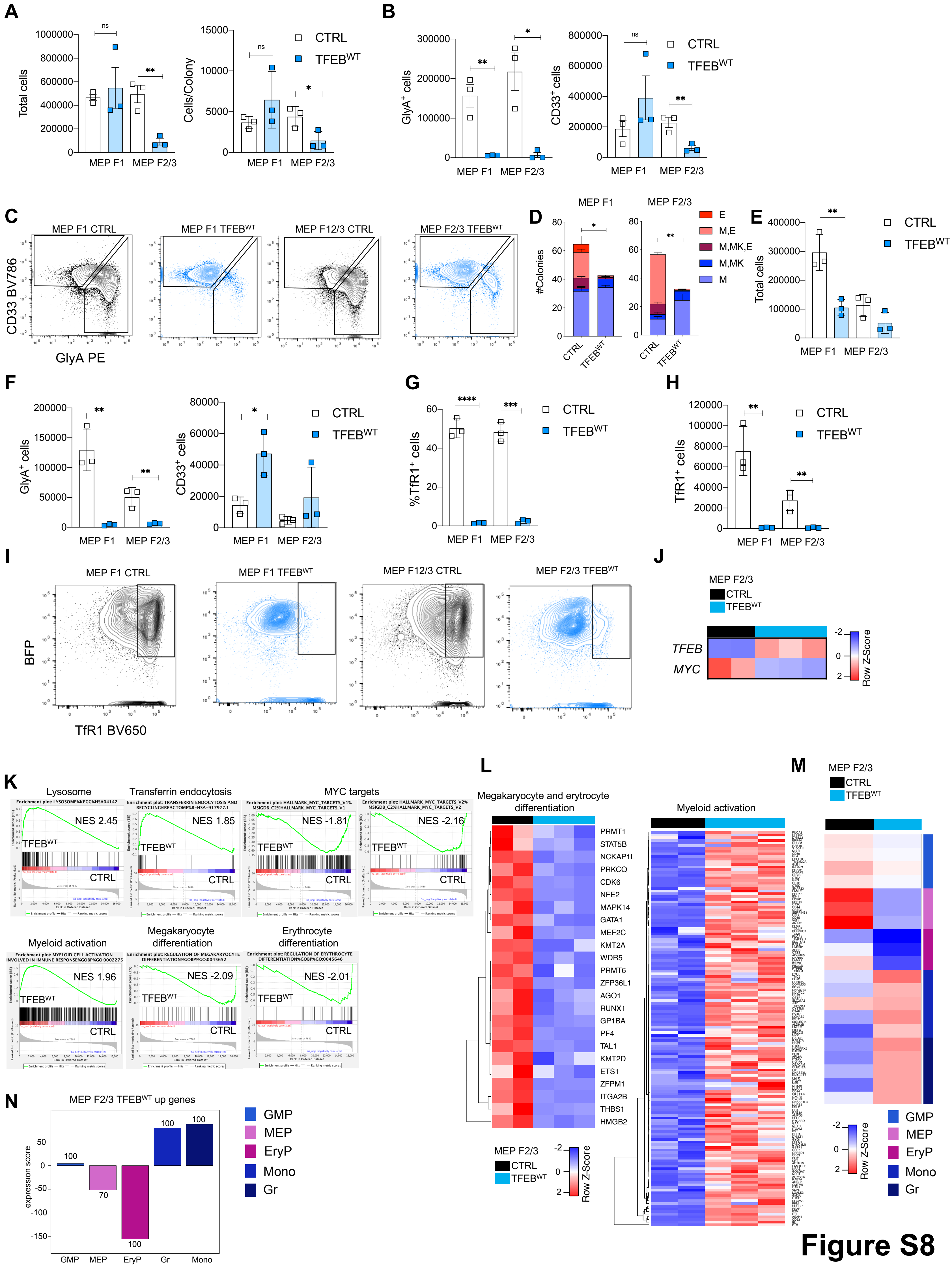
A
B
C
D
E
F
G
H
I
J
K
L
M
N
CTRL
TFEB^WT
MEP F1
MEP F2/3
Total cells
Cells/Colony
GlyA+ cells
CD33+ cells
MEP F1 CTRL
MEP F1 TFEB^WT
MEP F12/3 CTRL
MEP F2/3 TFEB^WT
CD33 BV786
GlyA PE
#Colonies
E
M,E
M,MK,E
M,MK
M
%TfR1+ cells
TfR1+ cells
BFP
TfR1 BV650
TFEB
MYC
Row Z-Score
Lysosome
Transferrin endocytosis
MYC targets
NES 2.45
NES 1.85
NES -1.81
NES -2.16
Myeloid activation
Megakaryocyte differentiation
Erythrocyte differentiation
NES 1.96
NES -2.09
NES -2.01
Megakaryocyte and erytrocyte differentiation
PRMT1
STAT5B
NCKAP1L
PRKCQ
CDK6
NFE2
MAPK14
GATA1
MEF2C
KMT2A
WDR5
PRMT6
ZFP36L1
AGO1
RUNX1
GP1BA
PF4
TAL1
KMT2D
ETS1
ZFPM1
ITGA2B
THBS1
HMGB2
MEP F2/3 TFEB^WT up genes
expression score
GMP
MEP
EryP
Mono
Gr
Figure S8

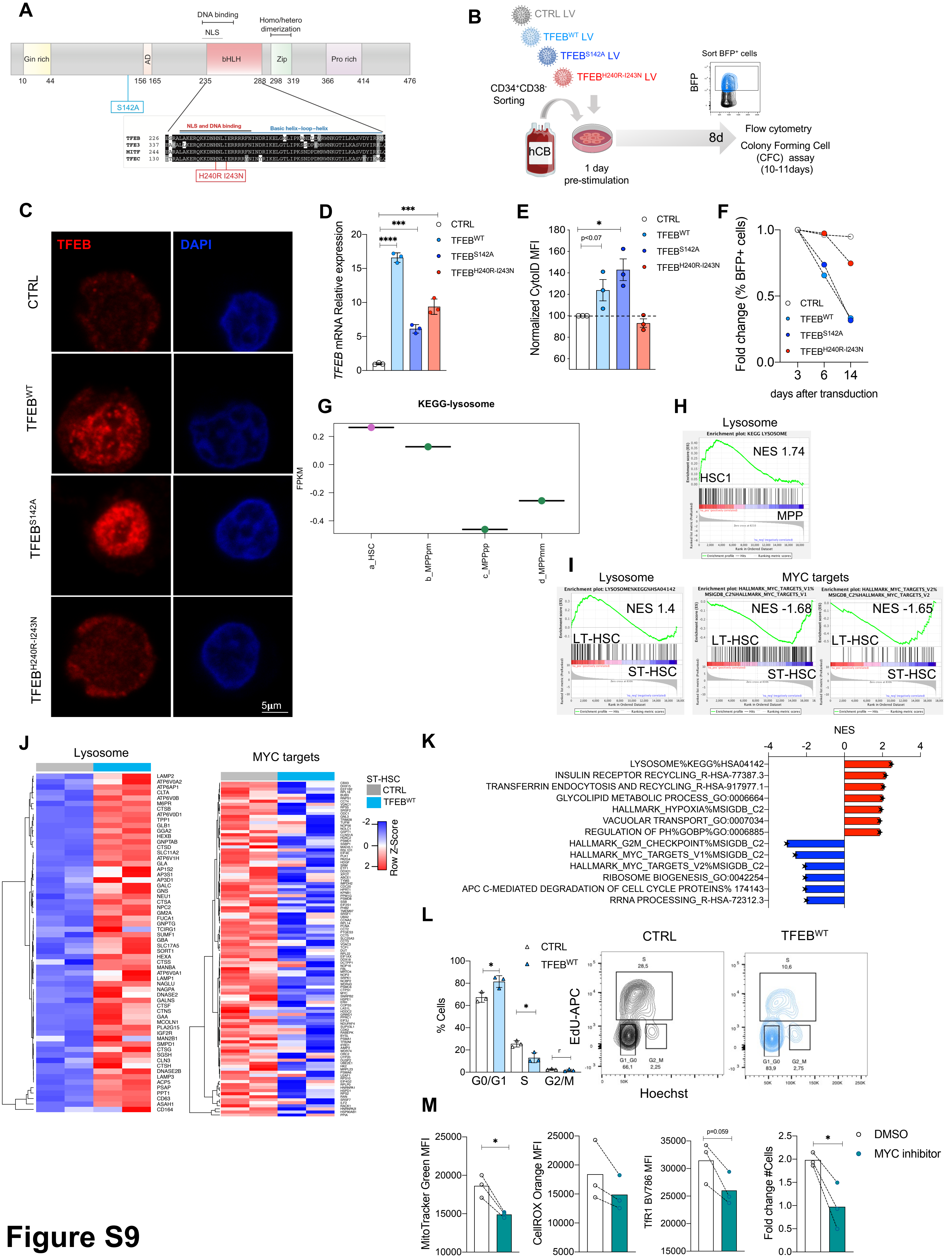
A
DNA binding
NLS
Homo/hetero dimerization
Gln rich
AD
bHLH
Zip
Pro rich
10 44 156 165 235 288 298 319 366 414 476
S142A
NLS and DNA binding
Basic helix-loop-helix
H240R I243N
B
CTRL LV
TFEB^WT LV
TFEB^S142A LV
TFEB^H240R-I243N LV
CD34+CD38- Sorting
hCB
1 day pre-stimulation
Sort BFP+ cells
BFP
8d
Flow cytometry
Colony Forming Cell (CFC) assay (10-11days)
C
TFEB
DAPI
CTRL
TFEB^WT
TFEB^S142A
TFEB^H240R-I243N
5µm
D
TFEB mRNA Relative expression
E
Normalized CytoID MFI
F
Fold change (% BFP+ cells)
days after transduction
CTRL
TFEB^WT
TFEB^S142A
TFEB^H240R-I243N
G
KEGG-lysosome
FPKM
a_HSC
b_MPPpm
c_MPPpp
d_MPPmm
H
Lysosome
NES 1.74
HSC1
MPP
I
Lysosome
MYC targets
NES 1.4
NES -1.68
NES -1.65
LT-HSC
ST-HSC
J
Lysosome
MYC targets
ST-HSC
CTRL
TFEB^WT
Row Z-Score
K
NES
LYSOSOME%KEGG%HSA04142
INSULIN RECEPTOR RECYCLING_R-HSA-77387.3
TRANSFERRIN ENDOCYTOSIS AND RECYCLING_R-HSA-917977.1
GLYCOLIPID METABOLIC PROCESS_GO:0006664
HALLMARK_HYPOXIA%MSIGDB_C2
VACUOLAR TRANSPORT_GO:0007034
REGULATION OF PH%GOBP%GO:0006885
HALLMARK_G2M_CHECKPOINT%MSIGDB_C2
HALLMARK_MYC_TARGETS_V1%MSIGDB_C2
HALLMARK_MYC_TARGETS_V2%MSIGDB_C2
RIBOSOME BIOGENESIS_GO:0042254
APC C-MEDIATED DEGRADATION OF CELL CYCLE PROTEINS% 174143
RRNA PROCESSING_R-HSA-72312.3
L
CTRL
TFEB^WT
% Cells
G0/G1
S
G2/M
EdU-APC
Hoechst
M
MitoTracker Green MFI
CellROX Orange MFI
TfR1 BV786 MFI
Fold change #Cells
DMSO
MYC inhibitor
p=0.059

### Figure S9

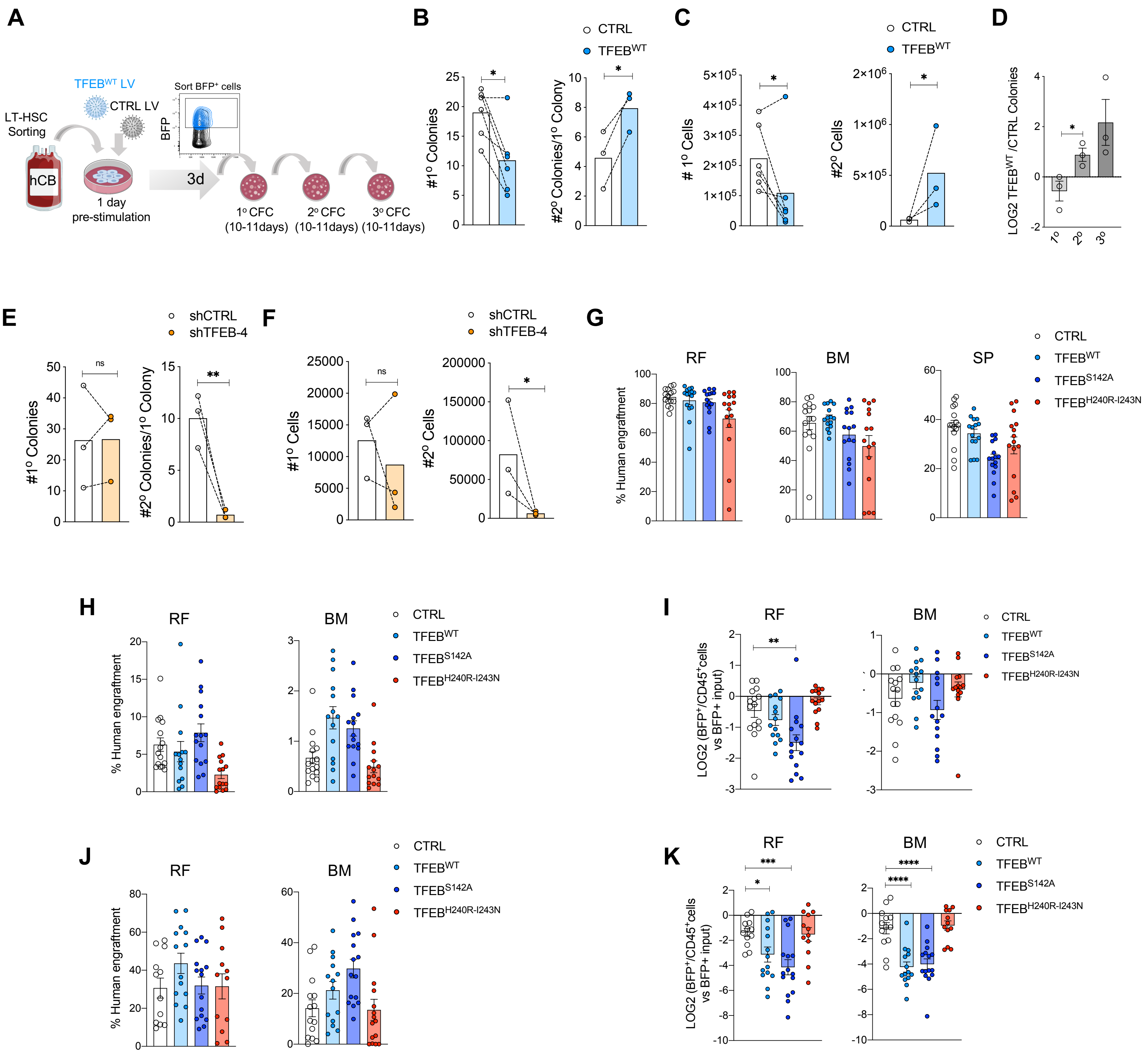


L

| | Cell doses | | | |
|---|---|---|---|---|
| | 40K | 75K | 150K | 250K |
| CTRL | 1/9 | 0/3 | 0/4 | 3/8 |
| TFEB$^{WT}$ | 3/9 | 2/2 | 2/4 | 2/2 |
| TFEB$^{S142A}$ | 5/7 | - | 2/2 | - |
| TFEB$^{H240R-I243N}$ | 0/8 | - | 3/5 | 0/2 |
| | Number of engrafted/injected mice | | | |

M

| | Stem cell frequency | | | | |
|---|---|---|---|---|---|
| | Lower | Estimate | Upper | Fold Increase | Pr(>Chisq) |
| CTRL | 1845579 | 691827 | 259336 | | |
| TFEB$^{WT}$ | 212178 | 103960 | 50937 | 6.65 | 6.60E-06 |
| TFEB$^{S142A}$ | 76703 | 31353 | 12815 | 22.06 | 0.00122 |
| TFEB$^{H240R-I243N}$ | 1391133 | 444120 | 141785 | 1.55 | 0.569 |

### Figure S10

A

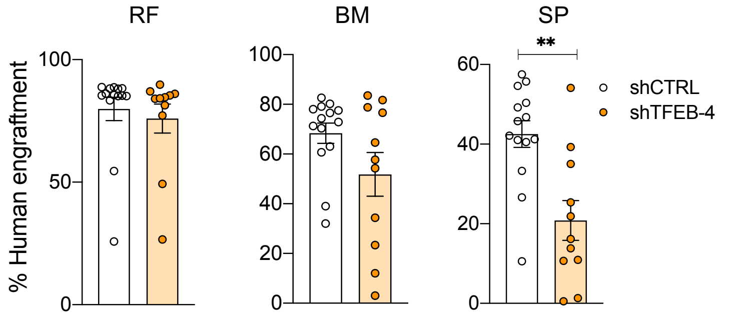


B

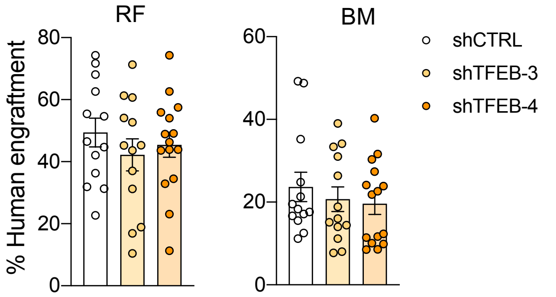


C

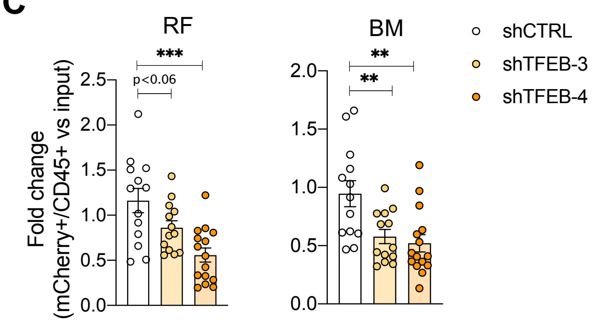


D

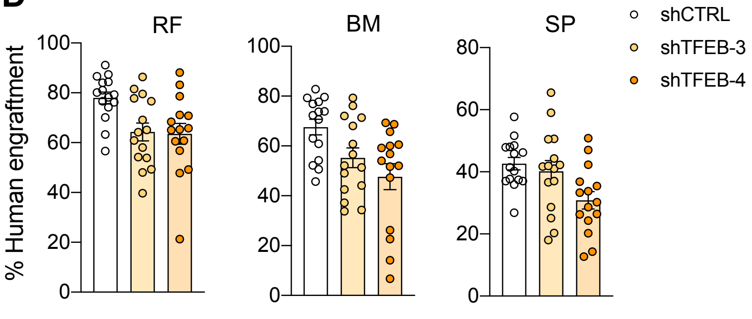


E

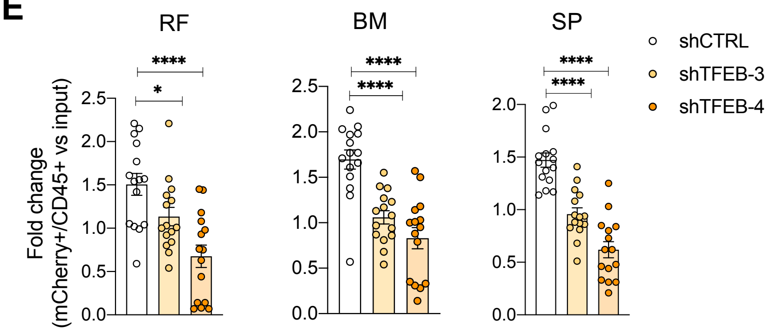


F

| | Cell doses | | |
|---|---|---|---|
| | 50k | 150k | 250k |
| shCTRL | 3/7 | 3/6 | 7/10 |
| shTFEB-4 | 0/8 | 1/6 | 5/10 |
| | Number of engrafted/injected mice | | |

G

| | Stem cell frequency | | | |
|---|---|---|---|---|
| | Lower | Estimate | Upper | Fold Increase |
| shCTRL | 323218 | 180366 | 100651 | |
| shTFEB-4 | 1123330 | 507535 | 229311 | -2.81 |

H

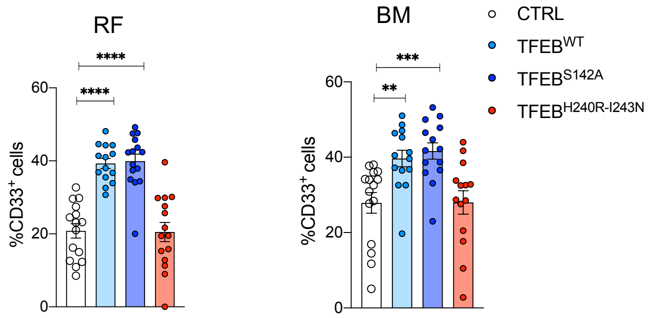


I

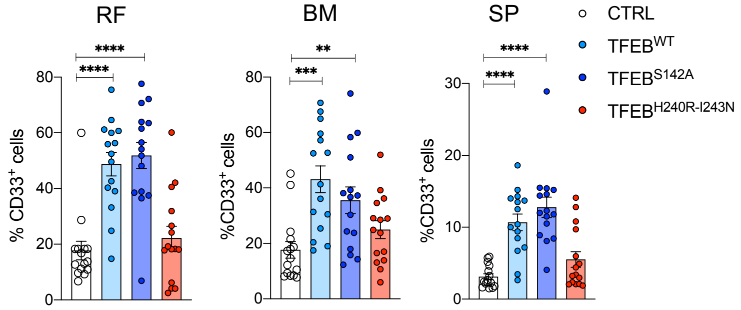


J

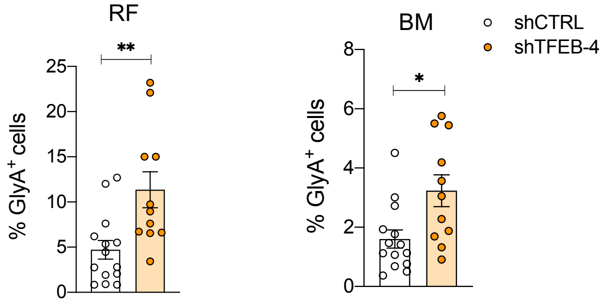


### Figure S11

A

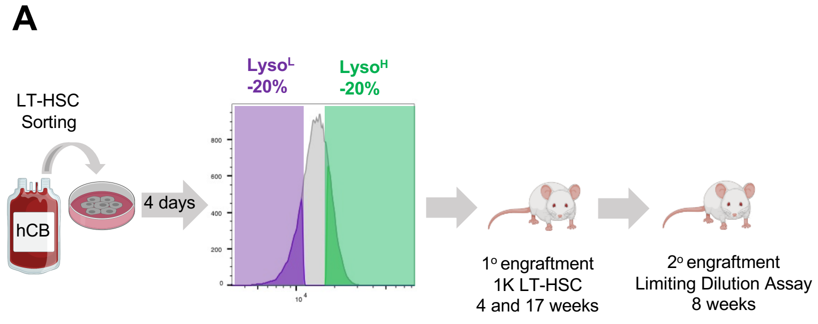


B

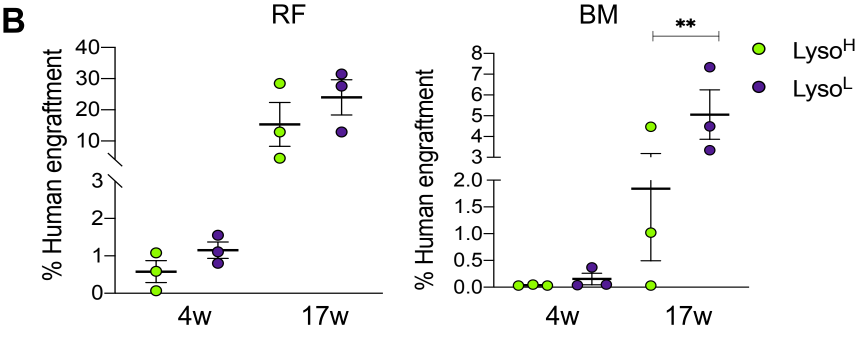


C

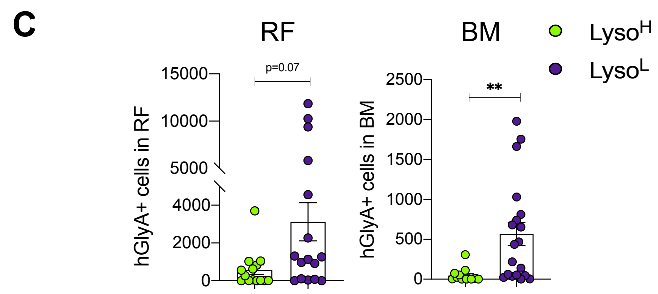


D

| | Cell doses | | |
|---|---|---|---|
| | 50k | 150k | 250k |
| LysoL | 1/17 | 1/11 | 4/15 |
| LysoH | 2/13 | 4/7 | 4/7 |
| | Number of engrafted/injected mice | | |

E

| | Stem cell frequency | | | |
|---|---|---|---|---|
| | Lower | Estimate | Upper | Fold Increase |
| LysoL | 469189.9 | 248591 | 131711.1 | |
| LysoH | 2087619 | 937599.2 | 421098.1 | 3.771653841 |

F

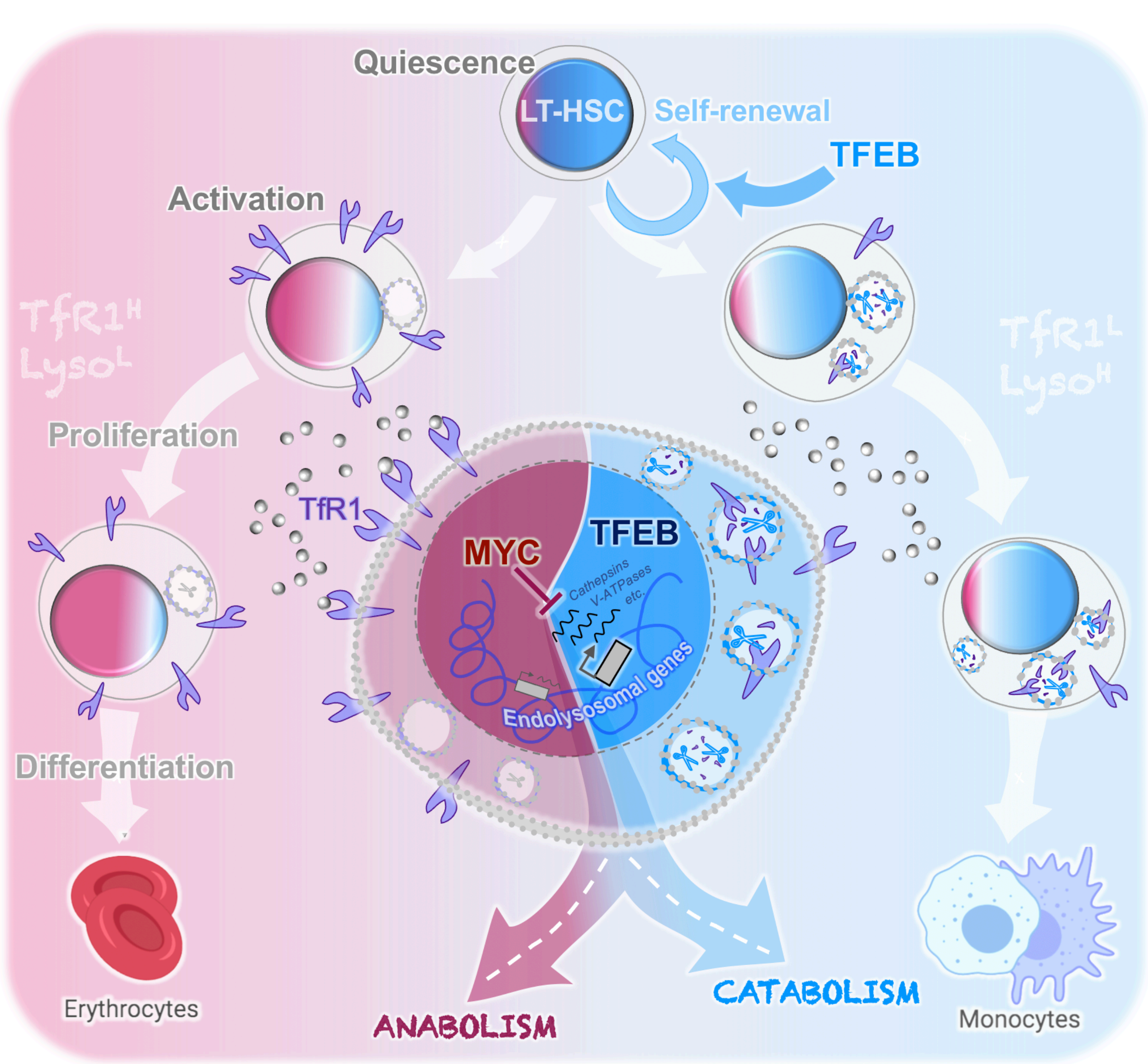


Figure S12
